# Activity profiling of deubiquitinating inhibitors-bound to SARS-CoV-2 papain like protease with antiviral efficacy in murine infection model

**DOI:** 10.1101/2022.11.11.516107

**Authors:** Shweta Choudhary, Sanketkumar Nehul, Santhosh Kambaiah Nagaraj, Rohan Narayan, Shalja Verma, Swati Sharma, Annu Kumari, Ruchi Rani, Ankita Saha, Debabrata Sircar, Abinaya Kaliappan, Shashank Tripathi, Gaurav Kumar Sharma, Pravindra Kumar, Shailly Tomar

## Abstract

SARS-CoV-2 papain-like protease (PLpro) is a key antiviral target as it plays a dual role in viral replication and in modulation of innate immune responses by deubiquitinating or deISGylating host proteins. Thus, therapeutic targeting of PLpro serves as a two-pronged approach to abate SARS-CoV-2. Interestingly, PLpro shares structural and functional similarities with the cellular deubiquitinating enzymes (DUBs) and in this study this fact has been exploited to identify DUBs inhibitors that target the Ubiquitin/ISG15 binding site and the known catalytic substrate binding pocket of PLpro. Among these identified compounds, flupenthixol, lithocholic acid, teneligliptin, and linagliptin markedly inhibited the proteolytic activity of purified PLpro and demonstrated potent antiviral efficacies against SARS-CoV-2 infection in a dose dependent manner. Treatment with lithocholic acid and linagliptin suppressed the expression levels of inflammatory mediators, thereby, restoring immune responses. Crystal structures of SARS-CoV-2 PLpro in complex with linagliptin and with lithocholic acid determined in this study, revealed insights into the inhibition mechanism with unique interactions within the Ubiquitin/ISG15 binding site (S2 site; Phe69, His73, Asn128, His175) and the substrate binding cleft. Additionally, oral and intraperitoneal treatments with linagliptin increased survival, reduced lung viral load, and ameliorated histopathological damage in mouse-adapted model of SARS-CoV-2 infection. The study for the first time demonstrates a two-pronged strategy using DUB inhibitors that target the proteolytic activity of PLpro and simultaneously reinstates the host’s immune response against SARS-CoV-2.

## Introduction

The COVID-19 pandemic, caused by the SARS-CoV-2 and the resurgence of variants of concern along with the drug-resistant variants, has taken a heavy toll on global health and calls for effective anti-SARS-CoV-2 drugs (Hirotsu et al., 2023; Iketani et al., 2023; Stevens et al., 2024). The rapid rollout of vaccine-based prophylactic therapies against SARS-CoV-2 initially saved numerous lives but the resulting immune pressures from vaccinations and infections led to emergence of highly contagious immune escape variants (Baker et al., 2021; Dubey et al., 2021; Mudgal et al., 2020). Concerns have been raised about the diminishing range, strength, and duration of vaccinal as well as natural immunity, which could compromise the geographical or temporal efficacy of these interventions and could lead to future breakthrough infections (Baker et al., 2021; Garcia-Beltran et al., 2021).

Precedent studies on SARS-CoV-2 highlight that the catalytic site of certain non-structural proteins (nsP) of SARS-CoV-2, such as papain-like protease (PLpro), 3-chymotrypsin-like protease (3CL pro or Mpro), RNA-dependent RNA polymerase (RdRp), and helicase display high levels of sequence similarity with their corresponding SARS and MERS counterparts and represent promising druggable targets for therapeutic development (Abdelnabi et al., 2021; Driouich et al., 2021; Li and De Clercq, 2020; Rani et al., 2022, 2020; Shan et al., 2021). Several small molecule therapeutics or neutralizing antibodies, either clinically approved or in late-stage of development, are targeting the key viral proteins such as the Spike protein, nucleocapsid, Mpro, and the RdRp (Choudhary et al., 2020; Cui et al., 2024; Dhaka et al., 2023; Krammer, 2020; Li et al., 2022; Singh et al., 2024); such as the FDA approved direct-acting antivirals, remdesivir and molnupiravir, which target the SARS-CoV-2 RdRp, and the Mpro inhibitor, Paxlovid (Kokic et al., 2021; Qiao et al., 2021; Wenhao et al., 2020; Zhang et al., 2020). Nevertheless, mutant SARS-CoV-2 strains with resistance against remdesivir and Paxlovid have been detected in cell culture and drug treated COVID-19 patients, raising a demand for additional antivirals with alternative modes of action (Hirotsu et al., 2023; Hu et al., 2023; Iketani et al., 2023; Stevens et al., 2024). Indeed, PLpro presents to be a more challenging and potentially promising druggable target of SARS-CoV-2 because of the shallow S1 and S2 sites, and effective antiviral drugs are yet to be developed for it (Tan et al., 2024).

The genome of SARS-CoV-2 comprises of a single-stranded positive-sense RNA of ∼30kb, which is translated to produce structural, non-structural, and accessory viral proteins, essential for carrying out replication, maturation, and genome packaging of SARS-CoV-2 (Arya et al., 2021; V’kovski et al., 2021). The genomic RNA gets translated to produce two overlapping polyproteins, pp1a and pp1ab (formed after −1 ribosomal frameshifting mechanism), that are proteolytically cleaved and processed into 16 nsPs by two virus-encoded cysteine proteases, the PLpro and the Mpro (Osipiuk et al., 2021; Ren et al., 2021). These nsPs form the replication and transcription complex (RTC) responsible for directing the process of transcription, replication, and maturation of the viral genome (Arya et al., 2021; V’kovski et al., 2021). SARS-CoV-2 PLpro, a domain within nsP3 protein, is a cysteine protease that recognizes the P4-P1 consensus LXGG sequence and cleaves the peptide bond between nsP1/nsP2 (LNGG↓AYTR), nsP2/nsP3 (LKGG↓APTK), and nsP3/nsP4 (LKGG↓KIVN) (Shan et al., 2021; Wioletta et al., 2021). It shares a sequence identity of 83% with PLpro of SARS-CoV and is relatively distant from MERS with a sequence identity of 32.9% (Shin et al., 2020; Weglarz-Tomczak et al., 2021). As an evasion mechanism against the host innate immune response, PLpro possesses deubiquitination and deISG15ylation activities that cleave off ubiquitin (Ub) and ISG15 (ubiquitin-like interferon-stimulated gene 15) post-translational modifications from host proteins by recognizing the C-terminal RLRGG sequence (McClain and Vabret, 2020; Shen et al., 2022; Shin et al., 2020). Interestingly, the multifunctional PLpro inactivates TBK1, a kinase required to activate transcription factor interferon responsive factor 3 (IRF3), prevents dimerization and translocation of IRF3 to the nucleus, and blocks NF-κB signaling pathway, eventually attenuating the type I interferon response in the host cell (Fu et al., 2021; McClain and Vabret, 2020; Osipiuk et al., 2021). Due to the dual role of PLpro in replication and pathogenesis of SARS-CoV-2, targeting PLpro may have an advantage in not only suppressing viral infection but also in restoring the dysregulated signaling cascades in infected cells that may otherwise lead to cell death of the surrounding non-infected cells. Despite extensive efforts in drug designing and lead optimization, the development of PLpro inhibitors is challenging and still remain at the preclinical stage with only a few covalent and non-covalent inhibitors reported till date (Choudhary et al., 2023; Fu et al., 2021; Kiira et al., 2008; Patchett et al., 2021).

Of note, PLpro has a right-handed “thumb-palm-finger” architecture which is similar to that of host deubiquitinating enzymes (DUBs), the ubiquitin-specific protease (USP), despite very low sequence similarity between them (∼10%) (Osipiuk et al., 2021). SARS-CoV-2 PLpro has two domains, the “thumb-palm-finger” catalytic domain at the C-terminus and a small N-terminal ubiquitin-like (UBL) domain (Fu et al., 2021; Klemm et al., 2020; Osipiuk et al., 2021). The active site of PLpro is made up of a canonical cysteine protease triad of Cys111, His272, and Asp286 residues, located at the interface of thumb and palm sub-domains. The most complex architecture is of the fingers subdomain, as it is made up of two α-helices, six β-strands, and a zinc-binding loop that is indispensable for maintaining structural integrity (Gao et al., 2021; Klemm et al., 2020; Osipiuk et al., 2021). An important flexible β-turn/loop, blocking/binding loop 2 (BL2 loop) that recognizes the P2-P4 of the LXGG motif of the substrate and closes upon binding of substrate/inhibitor, is located between *β*11–12 strands spanning the residues from 267–271 (Gao et al., 2021; Klemm et al., 2020; Shin et al., 2020). Interestingly, the PLpro shares high structural similarity and structural conservation of the active site architecture and orientation of the catalytic triad residues Cys-His-Asp/Asn with the human DUBs, and previous studies have reported few USP inhibitors against SARS-CoV and MERS (Chou et al., 2008).

DUB/USP inhibitors are classified into several types based on their molecular skeletons and the core functional groups (Oliveira et al., 2022): cyanopyrrolidine derivatives (Bashore et al., 2020a), thiophene derivatives (Chen et al., 2017), quinazoline derivatives (Lamberto et al., 2017), piperidine derivatives (Lamberto et al., 2017), 2-amino-4-ethylpyridin derivatives (Kategaya et al., 2017), and Thienopyridine derivatives (Leger et al., 2020). Moreover, inhibitors with a cyanopyrrolidine warhead, such as gliptins, are reported to initiate a β-elimination reaction in the active site of USPs, resulting in irreversible inactivation of active site cysteine to dehydroalanine after desulfhydration reactions (Bashore et al., 2020b; Jadav et al., 2012). These gliptins are already in clinical use as an antidiabetic compound inhibiting dipeptidyl peptidase 4 (DPP4), a serine protease and a possible receptor for SARS-CoV-2 entry (Fisman and Tenenbaum, 2015; Jadav et al., 2012; Pitocco et al., 2020). Notably, the DPP4 inhibitors, linagliptin and sitagliptin are reported to exert anti-inflammatory effects on the liver and the lungs by inhibiting the IL-1 and IL-6 inflammatory pathways respectively (Kawasaki et al., 2018; Valencia et al., 2020; Yang et al., 2021; Zhuge et al., 2016). Therefore, DUB/USP inhibitors as PLpro inhibitors are proposed to have a dual mechanism of action in inhibiting SARS-CoV-2. The first inhibition mechanism potentially targets the viral polyprotein processing enzymatic activity of PLpro due to structurally conserved active site fold, while second mechanism is via restoration of the host’s innate immune responses.

The present study uses DUBs/USPs dual inhibitors that target the PLpro of SARS-CoV-2 by engaging the PLpro catalytic active site and the ISG15/Ub S2-binding pocket and also restore the host’s innate immune responses: an approach which has not been previously exploited for development of SARS-CoV-2 therapeutics. Structure-based docking and simulation were performed for identification of DUBs/USPs inhibitors that fit into the target PLpro site. Five potential compounds with binding comparable to known PLpro inhibitor, GRL0617 were identified (Fu et al., 2021). Two of these compounds, lithocholic acid and linagliptin exhibited significant binding affinity, and greater inhibitory potency against purified PLpro enzyme and high efficacy against SARS-CoV-2 viral replication across multiple cell lines. Poly I:C stimulated cells treated with lithocholic acid and linagliptin demonstrate reduced expression levels of inflammatory mediators IL-6 and TNFα, supporting the restoration of a balanced immune response. Next, to dissect the mechanism of action of identified inhibitors, the crystal structures of linagliptin and lithocholic acid in complex with PLpro were determined to expose inhibitory sites and to gain structural insights into the inhibitory mechanisms. Further, the compound, linagliptin, was selected as an optimized lead and it’s anti-SARS-CoV-2 activity was characterized *in vivo* using a mouse-adapted SARS-CoV-2 model. Herein, this study opens a new avenue for developing DUBs inhibitor based dual-mechanistic antiviral therapeutics against SARS-CoV-2.

## Results

### PLpro: a virus-encoded DUB structural mimic

SARS-CoV-2 PLpro has two functions in the virus life cycle. As a viral cysteine protease, it plays an essential role in the cleavage and processing of the viral polyproteins, and as a viral deubiquitinase, it functions in the evasion of host innate immune responses by removing ISG15/ubiquitin from the host proteins. Structural studies in the last decade have contributed to an understanding of SARS-CoV-2 PLpro function as a deubiquitinating enzyme. We compared the structures of PLpro with the host USPs/DUBs by structure superimposition to investigate the structural similarity and mimicry between the PLpro and the host DUBs. A detailed structural comparison of the PLpro with UPS7 (PDB ID: 2F1Z) (Hu et al., 2006) and USP2 (PDB ID: 5XU8) (Chuang et al., 2018) proteins was carried out using the PDBeFold (SSM: Secondary Structure Matching) and multiple sequence alignment was performed with ESPript3 (Figure S1). Although these proteins share a very low percentage of sequence identity with an alignment of only 187 residues of PLpro with USP7 and USP2 proteins, the structural superimposition topologies were reasonably similar with an overall root mean square deviation (RMSD) value of 1.68Å and 1.24 Å for USP7 and USP2, respectively (Figure 1A-C). Reminiscent of the overall architecture of DUBs of the USP family, PLpro shared a similar active site architecture with the host USP7 (also known as HAUSP) and USP2 proteins, including the signature catalytic triad (Cys-His-Asp/Asn) at the interface of its thumb and palm domains. Briefly, the palm subdomain of PLpro, USP7, and USP2 comprises of 6 β-sheets and includes the conserved catalytic triad (Cys-His-Asp/Asn) of the active site (Figure 1A-C). Beyond the catalytic residues, USPs contain N- and C-terminal extensions that aid in substrate recognition and binding. The BL2 loop of SARS-CoV-2 PLpro (residues 267–272) that provides floor to ubiquitin tail, also superimposes well with the BL2 loops of USP7 and USP2 (Figure 1C) (Pozhidaeva et al., 2017; Wertz and Murray, 2019). Interestingly, the zinc-binding domain located in the thumb region of SARS-CoV-2 comprises of 4 cysteine residues (Cys189, Cys192, Cys224, and Cys226) and the same pattern is observed for USP2 as well (Cys334, Cys337, Cys381, and Cys384) (Renatus et al., 2006). Contrastingly, some of the cysteine residues of this metal binding motif have been mutated during evolution in USP7 and other DUBs due to which the zinc binding ability was lost while the fold integrity has still been retained (Renatus et al., 2006). Importantly, residues Gly269, Gly271, His272, and Tyr273 (numbering as per PLpro) are conserved between USP7 and USP2 indicating as expected that the substrate binding or proteolytic cleft formed near the BL2 loop are also conserved (Figure 1 and Figure S1) (Gao et al., 2021; Kiira et al., 2008). Nevertheless, the structural architecture of the catalytic sites are highly similar in PLpro, USP7, and USP2, suggesting that some of the DUBs/USPs inhibitors or their analogs may display inhibitory activity against PLpro proteases.

**Figure 1.**
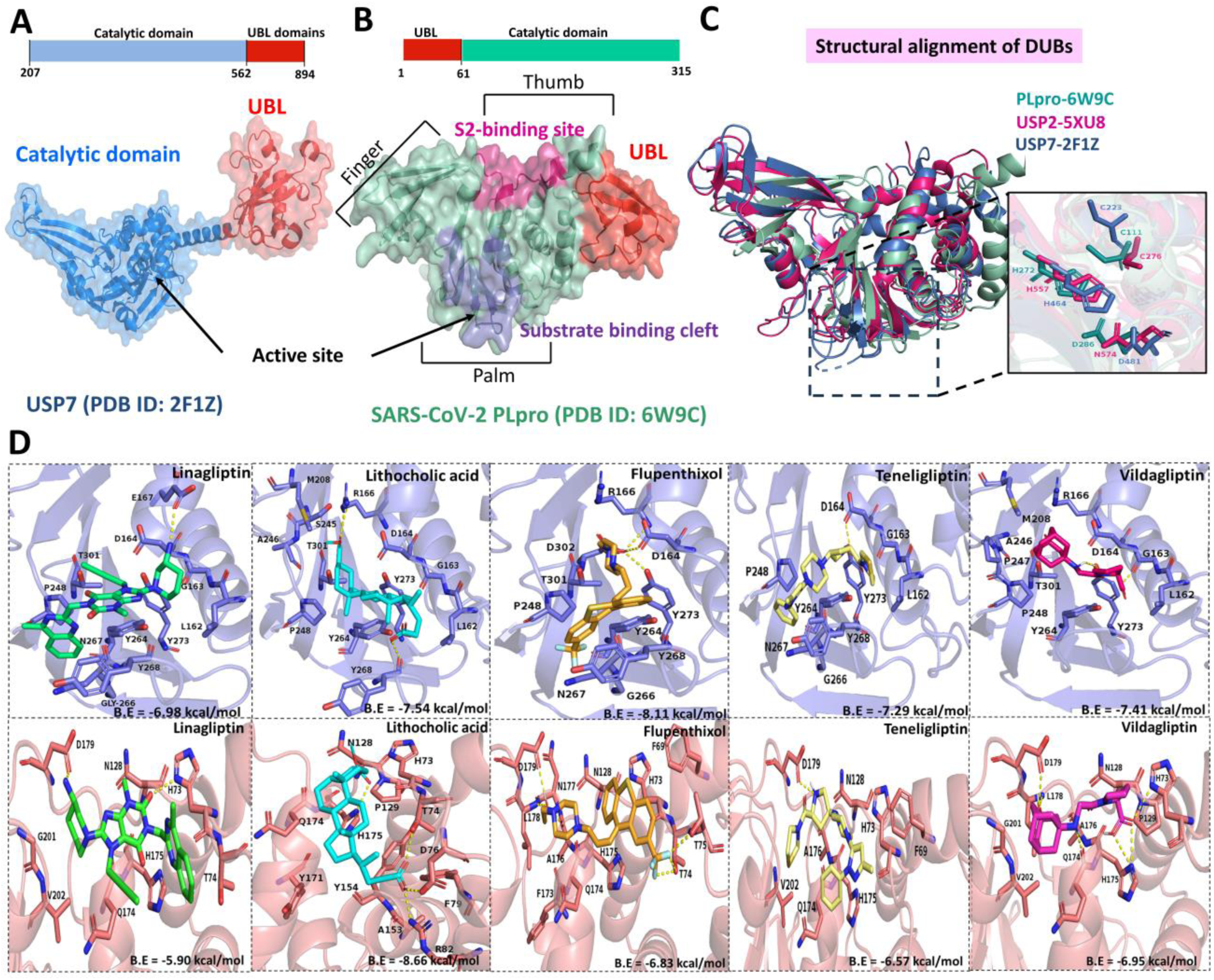
Domain organization and comparison of structural motifs of SARS-CoV-2 PLpro with the cellular DUBs. (A) Surface representation of the domain organization and structural architecture of USP7 (PDB ID: 2F1Z). The catalytic domain is represented in blue and the UBL domain is depicted as red surface. (B) Surface view representation of the overall structure of PLpro (PDB ID: 6W9C) with thumb, palm, and zinc-finger domains marked. The substrate binding cleft is depicted as violet surface, the S2 Ub/ISG15 binding site is represented as pink surface, and the UBL domain is shown as red surface. (C) Structural alignment of secondary structure elements of PLpro (Purple; PDB ID: 6W9C), with the cellular DUBs USP7 (Green; PDB ID: 2F1Z), and USP2 (Magenta; PDB ID: 5XU8). A total of 187 residues of each protein chosen by web-based server PDBeFold, were aligned based on the secondary structure elements. Cartoon representation shows PLpro in green, USP2 in pink, and USP7 in blue. The positioning of catalytic triad residues is represented in magnified view. Catalytic triad residues Cys, His, and Asp of PLpro, USP2, and USP7 represented in sticks, are completely superimposed. (D) PyMOL schematic representations demonstrating binding modes and key hydrophobic/hydrogen interactions of the top-hit screened ligands docked at the substrate-binding cleft (purple) and Ub/ISG15 S2-binding site (brick red) of PLpro.

### Identification of PLpro binding DUBs inhibitors

Crystal structures of PLpro in complex with GRL0617, HE9, and acriflavine inhibitors provides the basis for targeting the multiple sites on SARS-CoV-2 PLpro with DUBs inhibitors (Gao et al., 2021; Napolitano et al., 2022; Srinivasan et al., 2022). To achieve this, we made an *in-house in silico* library of small molecules containing DUB inhibitors including cyanopyrrolidine ring-containing compounds, quinazoline derivatives, and piperidine derivatives (Bashore et al., 2020a; Chen et al., 2017; Kategaya et al., 2017; Lamberto et al., 2017; Leger et al., 2020; Oliveira et al., 2022; Zhuge et al., 2016). This library of molecules was subjected to molecular docking using in-built AutoDock Vina module of PyRx 0.8 (Table S1). The GRL0617 (control inhibitor molecule) binding site in the PLpro substrate binding pocket or active site (residues Leu162, Gly163, Asp164, Glu167, Met208, Pro247, Pro248, Tyr264, Tyr268, Gln269, Gly271, and Tyr273, Thr301: PDB ID: 7CJM) was used for docking the DUBs inhibitors (Fu et al., 2021; Klemm et al., 2020; Yuan et al., 2022). Additionally, the molecules predicted to bind PLpro active site more effectively than GRL0617 were further docked into the ISG15 binding S2-binding site in the PLpro structure (PDB: 6YVA). The crystal structures of SARS-CoV-2 PLpro in complex with ISG15 (PDB: 6YVA) has a distal S2-binding site located on PLpro thumb domain, contributing to substrate recognition and differential cleavage (Gao et al., 2021; Shin et al., 2020). The key interacting residues of PLpro mediating contacts with the N-terminal ISG15 at the S2-binding site (Val66, Phe69, Glu70, Thr75, Asn128, Tyr171, Asn177, and Asp180) were used for docking the selected DUBs inhibitors (Patchett et al., 2021)(Riva et al., 2020). In conclusion, targeting the substrate-binding cleft and the S2 Ub/ISG15 binding sites of PLpro represents a new approach for preventing viral replication and its interaction with host cellular pathways.

A total of 20 compounds were shortlisted based on binding affinities with the substrate binding cleft and RMSD values (<1 Å) and the scores of these 20 compounds are listed in Table S2. Among these top hits, the compounds pimozide, MF094, FT671, ML323, FT827, GNE6640, and N-cyanopyrrolidine have already been recently reported as protease inhibitors of SARS-CoV-2 PLpro, thus validating our developed protocol (Cho et al., 2022; Shan et al., 2021). Additionally, the published literature highlights efficacy of sitagliptin and gemigliptin in type II diabetes mellitus patients infected with SARS-CoV-2 when administered as a monotherapy or in the form of a combination therapy (Alhakamy et al., 2021; Hazra, 2021; Narayanan et al., 2022). Therefore, these compounds were not utilized for further studies. The remaining compounds of the top list were further investigated for their binding modes and stabilities of protein-ligand complexes. Based on binding energies, number of polar/hydrophobic interactions with the substrate binding site, and commercial availability, a total of five compounds were selected for detailed analysis by molecular docking into the catalytic substrate-binding site and S2-binding site and simulation analysis using AutoDock4.2.6 and Gromacs. The binding affinities of the top-screened compounds at the catalytic substrate-binding site ranged between −6.9 kcal/mol and −8.1 kcal/mol (Table 1), which are higher or similar to the docking score for the control inhibitor ligand GRL0617 (−6.94 kcal/mol). Flupenthixol compound displayed highest binding affinity of −8.11 kcal/mol for substrate-binding site, forms four hydrogen-bonds (H-bond) with active site residues Asp164, Arg166, Tyr273, and Asp302 (Figure 1D, Figure S2, Table 1) and additional hydrophobic interactions with PLpro residues Pro248, Tyr264, Gly266, Asn267, Tyr268, Thr301 that contribute to the stabilization of protein-ligand complex (Figure 1D, Table 1). Lithocholic acid was the second-best candidate that forms three H-bonds with targeted amino acids of catalytic site, Arg166 and Tyr268, along with ten hydrophobic interactions (Figure 1D and Table 1). Like other top hits, teneligliptin and linagliptin also occupied catalytic substrate-binding site of PLpro forming H-bonds with key GRL0617 binding residues (Asp164: teneligliptin; Glu167: linagliptin) (Figure 1D, Table 1, and Figure S2). Teneligliptin and linagliptin protein-ligand complexes are further stabilized by extensive hydrophobic interactions primarily involving Leu162, Gly163, Pro248, Tyr264, Gly266, Asn267, and Tyr268 substrate binding residues of PLpro (Figure 1D, Table 1, and Figure S2). Similarly, vildagliptin also displayed a network of two H-bonds and ten hydrophobic interactions with the targeted site (Figure 1D, Figure S2, and Table 1). Molecular docking studies at the S2-Ub/ISG15 binding site of PLpro revealed binding affinities in the range of −5.9 to −8.6 kcal/mol. Lithocholic acid, displayed highest binding energy at the S2-binding site making molecular contacts with His73, Thr74, and Phe79, the key substrate interacting residues of S2 site (Figure 1D, Table 1). Likewise, linagliptin interacted with His73 and Asp179 through H-bond interactions along with additional stabilizing hydrophobic interactions with key residues of S2 site (Figure 1D, Table 1). The compound flupenthixol sustained seven H-bond interactions and other additional hydrophobic interactions (Figure 1D, Table 1). PLpro-teneligliptin involved one H-bond with Asp179 and made additional hydrophobic interactions at the S2 site (Table 1). The binding of compound vildagliptin at S2 site was stabilized by six H-bond and additional hydrophobic interactions, as shown in Table 1 (Figure 1D, Figure S2). The stability of these protein-ligand complexes were further assessed by performing molecular dynamics (MD) simulation for 100 ns (Figure S3). The complexes attained equilibrium after 20 ns and were stable throughout the run. The overall ligand/protein complexes showed no major differences in the overall stability compared to native PLpro, with an average RMSD value in the range of 0.1-0.4 Å for all complexes (Figure S3).

**Table 1.** Binding energies, hydrogen, and hydrophobic interactions of the selected PLpro of SARS-CoV-2 inhibitors with the key residues of the catalytic site and Ub/ISG15 S2-binding site.

| Name of compound | Site | AutoDock Binding energies (kcal/mol) | Hydrogen bonds |  | Hydrophobic interactions |
| --- | --- | --- | --- | --- | --- |
|  |  |  | Bonds (lig - protein) | Bond length (Å) |  |
| Flupenthixol | Catalytic-substrate binding site | -8.11 | O-OD1 (Asp164) | 3.17 | Pro248, Tyr264, Gly266, Asn267, Tyr268, and Thr301 |
|  |  |  | O-N (Arg166) | 3.16 |  |
|  |  |  | O-OH (Tyr273) | 2.95 |  |
|  |  |  | O-OD2 (Asp302) | 2.71 |  |
|  | Ub/ISG15 S2-binding site | -6.83 | F1-OG1 and F2-OG1 (Thr74) | 2.69 and 2.6 | Phe69, His73, Asn128, Phe173, Gln174, His175, Asn177 |
|  |  |  | F2-N and F2-OG1 (Thr75) | 1.10 and 3.12 |  |
|  |  |  | O-O (Ala176) | 2.67 |  |
|  |  |  | O-N (Leu178) | 2.98 |  |
|  |  |  | O-OD1(Asp179) | 2.75 |  |
| Lithocholic acid | Catalytic-substrate binding site | -7.54 | O4-NE & O3-NH2 (Arg166) | 2.87 and 3.05 | Leu162, Gly163, Asp164, Met208, Ser245, Ala246, Pro248, Tyr264, Tyr273, and Thr301 |
|  |  |  | O2-O (Tyr268) | 2.80 |  |
|  | Ub/ISG15 S2-binding site | -8.66 | O1-OG1(Thr74) | 2.81 | His73, Phe79, Asn128, Pro129, Ala153, Tyr154, Tyr171, Gln174 |
|  |  |  | O2-OD2(Asp76) | 2.86 |  |
|  |  |  | O2-NH2(Arg82) | 2.73 |  |
|  |  |  | O3-ND1 (His175) | 2.72 |  |
| Vildagliptin | Catalytic-substrate binding site | -7.41 | O4-O (Gly163) | 2.55 | Leu162, Asp164, Arg166, Met208, Ala246, Pro247, Pro248, Tyr264, and Thr301 |
|  |  |  | N1-OH and O2-OH (Tyr273) | 3.19 and 3.09 |  |
|  | Ub/ISG15 S2-binding site | -6.95 | O3-ND1 (His73) | 3.14 | Asn128, Pro129, Ala176, Gly201, Val202 |
|  |  |  | N1-O (Gln174) | 2.74 |  |
|  |  |  | O3-ND1 and O4-ND1 (His175) | 2.98 and 3.16 |  |
|  |  |  | O1-N (Leu178) | 2.84 |  |
|  |  |  | C4-OD1(Asp179) | 2.71 |  |
| Teneligliptin | Catalytic-substrate binding site | -7.29 | N3-OD1 (Asp 164) | 2.41 | Leu162, Gly163, Pro248, Tyr264, Gly266, Asn267, and Tyr268 |
|  |  |  | N3-OH (Tyr 273) | 2.78 |  |
|  | Ub/ISG15 S2-binding site | -6.57 | N3-OD1(Asp179) | 2.58 | Phe69, His73, Asn128, Gln174, His175, Ala176, Val202 |
| Linagliptin | Catalytic-substrate binding site | -6.98 | N27-OE1 (Glu167) | 3.11 | Leu162, Gly163, Asp164, Pro248, Tyr264, Gly266, Asn267, Tyr268, Tyr273, and Thr301 |
|  |  | -5.90 | O19-ND1 (His73) | 3.03 |  |
|  | Ub/ISG15<br>S2-binding<br>site |  | N27-OD1 (Asp179) | 2.59 | Thr74, Asn128,<br>Gln174, His175,<br>Gly201, Val202 |

### Inhibition of PLpro protease activity

A fluorescence resonance energy transfer (FRET)-based protease assay utilizing the Z-RLRGG-AMC fluorescent substrate, was employed to evaluate the inhibitory efficacy of selected compounds against PLpro. SARS-CoV-2 PLpro and its inactive mutant (PLproC111S) were expressed in *Rosetta* DE3 and subsequently purified using affinity chromatography. A single band at ∼36 kDa in sodium dodecyl sulphate-polyacrylamide gel electrophoresis (SDS-PAGE) confirmed PLpro protein purity (Figure 2A). The enzymatic activity of purified PLpro was validated through a FRET assay which demonstrated that fluorescence signal production was exclusively associated with the proteolytic function of the PLpro, and not with the inactive PLproC111S mutant (Figure 2A and 2B). The same FRET assay was used to assess the inhibitory efficacy of the identified compounds, flupenthixol, lithocholic acid, vildagliptin, teneligliptin, and linagliptin against the enzymatic activity of PLpro. Among these selected compounds, both lithocholic acid and linagliptin robustly inhibited proteolytic activity of PLpro with > 80% inhibition of SARS-CoV-2 PLpro and half-maximal inhibitory concentration (IC_50_) values of 27.5 ± 2.20 μM and 53.1 ± 0.01 μM, respectively (Figure 2D and 2E). The compounds flupenthixol and teneligliptin showed IC_50_ values of 53.5 ± 4.09 μM and 259.2 ± 0.72 μM (Figure 2F and 2G). However, vildagliptin did not show any significant inhibition and has a maximum inhibitory effect of less than 50 percent only (IC_50_ > 500 μM) (Figure 2C). Because of low inhibitory potential of vildagliptin, it was not investigated further for binding kinetic assays using Surface Plasmon Resonance (SPR) and cell-based antiviral studies. Further, remaining selected four compounds were subjected to a series of SPR-based binding assays and cell-based *in vitro* studies.

**Figure 2.**
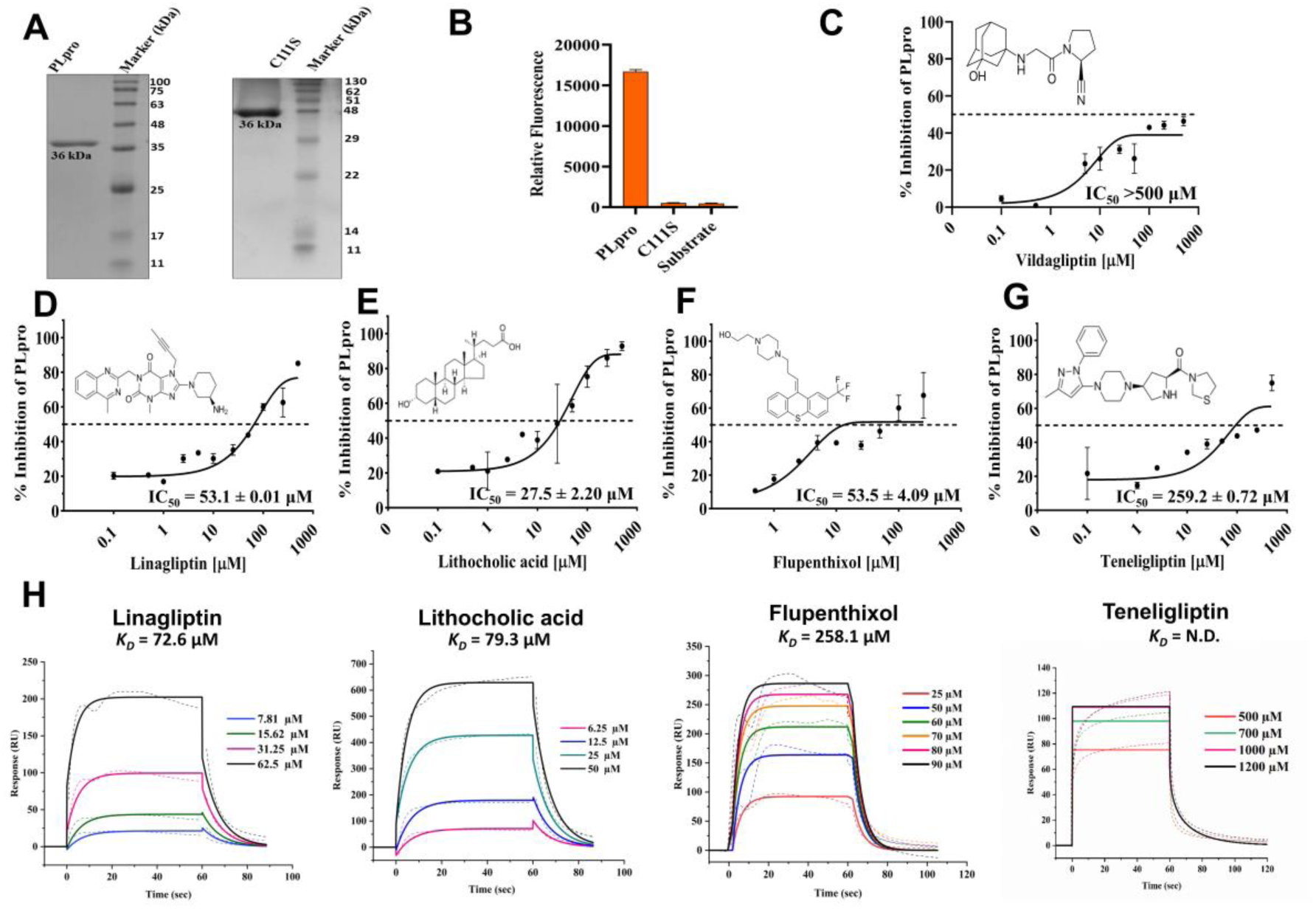
Binding affinity analysis and inhibition of PLpro protease activity. (A) SDS-PAGE profile of purified PLpro (∼36 kDa) and its inactive mutant C111S (B) FRET based protease activity comparison of wild type and C111S mutant (inactive) PLpro. Inhibition of proteolytic activity of PLpro by selected compounds. The inhibitory potential of identified compounds (C) vildagliptin, (D) linagliptin, (E) lithocholic acid, (F) flupenthixol, and (G) teneligliptin against PLpro were tested using fluorogenic peptide Z-RLRGG-AMC as the substrate. Black dotted lines indicate the concentration of compound needed to inhibit the proteolytic activity of PLpro by half (IC_50_). All data are shown as mean ± SEM, n = 2 independent experiments (H) Evaluation of binding affinities of lithocholic acid, linagliptin, flupenthixol, and teneligliptin to SARS-CoV-2 PLpro using SPR. Sensorgrams elucidating the binding kinetics of selected analytes injected successively at increased concentrations against immobilized SARS-CoV-2 PLpro. The data was fitted using 1:1 Langmuir binding model. The processed sensograms are color coded, and the binding response is increasing with the increase in the concentration of the analyte. K_D_, dissociation constant; RU, response units.

### Binding affinities of DUB inhibitors to PLpro

Prioritized lead compounds were tested in SPR assay to examine the binding kinetics and affinity of compounds against purified PLpro. After immobilization of histidine-tagged PLpro on Ni-nitrilotriacetic acid (Ni-NTA) chip, serial dose of selected compounds was allowed to flow over the chip and the time dependent optical signal was recorded. Consistent with the results of FRET assay, a dose-dependent increase in response was observed for binding of lithocholic acid, linagliptin, and flupenthixol to SARS-CoV-2 PLpro. As represented in Figure 2H, the *K_D_* values determined by SPR measurements for lithocholic acid-PLpro, linagliptin-PLpro, and flupenthixol-PLpro are 79.3 μM, 72.6 μM, and 258.1 μM, respectively. SPR data suggested that lithocholic acid and linagliptin bind to PLpro with stronger affinity in comparison flupenthixol (Figure 2H). While fitting the data for teneligliptin, the derivatives of the binding response and the dissociation were nonlinear and attempts to fit the data with 1:1 Langmuir Binding model failed with teneligliptin (data not shown). Together with results of fluorescence assay, SPR data confirms that these top-hits can be taken up for cell-based *in vitro* studies.

### Inhibition of SARS-CoV-2

The antiviral efficacy of selected compounds was investigated using *in vitro* cell based assays. First, to determine the non-cytotoxic concentrations of compounds, uninfected Vero cells, HEK-293T-ACE2, and A549-ACE2 were incubated with increased concentrations of compounds and cytotoxicity was measured using colorimetric [3-(4,5-dimethylthiazol-2-yl)-2,5-diphenyl tetrazolium bromide] (MTT) assay. Lithocholic acid, linagliptin, and teneligliptin exerted minimal cytotoxic effects across all cell lines at concentrations up to 50 μM (Figure S4). However, flupenthixol displayed cytotoxicity at concentrations above 50 μM in the tested cell lines (Figure S4). Half-maximal cytotoxicity (CC_50_) values were calculated and the testing of compounds in antiviral assays was performed at concentrations below CC_50_ values. The *in vitro* antiviral efficacy against clinical isolate of SARS-CoV-2 was investigated using Vero cells. The cell lysates of treated and untreated infected Vero cells were harvested to determine the viral load for quantifying the levels of viral RNA by real-time quantitative PCR (qRT-PCR) using SAR-CoV-2 RdRp and N-protein specific primers. Consistent with the results of biophysical studies, lithocholic acid and linagliptin effectively inhibited replication of SARS-CoV-2 in Vero cells, exhibiting an EC_50_ value of ∼21 μM and ∼14.65 μM, respectively (Figure 3). Interestingly, flupenthixol, which was observed to be a relatively weak inhibitor of PLpro enzymatic activity, also displayed inhibitory effect against SARS-CoV-2 at its non-toxic concentrations with an EC_50_ value of ∼5.10 µΜ and teneligliptin inhibited SARS-CoV-2 infection with EC_50_ ∼20.82 μM (Figure 3). TCID_50_ studies were carried out to validate qRT-PCR results and effect of compounds on cells infected with SARS-CoV-2 was assessed by performing the virus titration. The sharp decrease in viral titer was observed as compared to virus infected untreated positive control for linagliptin (decrease of ∼4 log TCID_50_/mL) and teneligliptin (decrease of ∼2.5 log TCID_50_/mL) (Figure S5). Flupenthixol at 20 µM concentration decreased the viral titer by ∼1.5 log TCID_50_/mL and 50 µM lithocholic acid reduced the titer by ∼1 log TCID_50_/mL as compared to virus control (Figure S5). Thus, treatment of cells with compounds significantly reduced SARS-CoV2 infectious virus production and cell death associated with virus infection. Together, the data suggest that these molecules displayed a dose-dependent inhibition of the virus replication in Vero cells.

**Figure 3.**
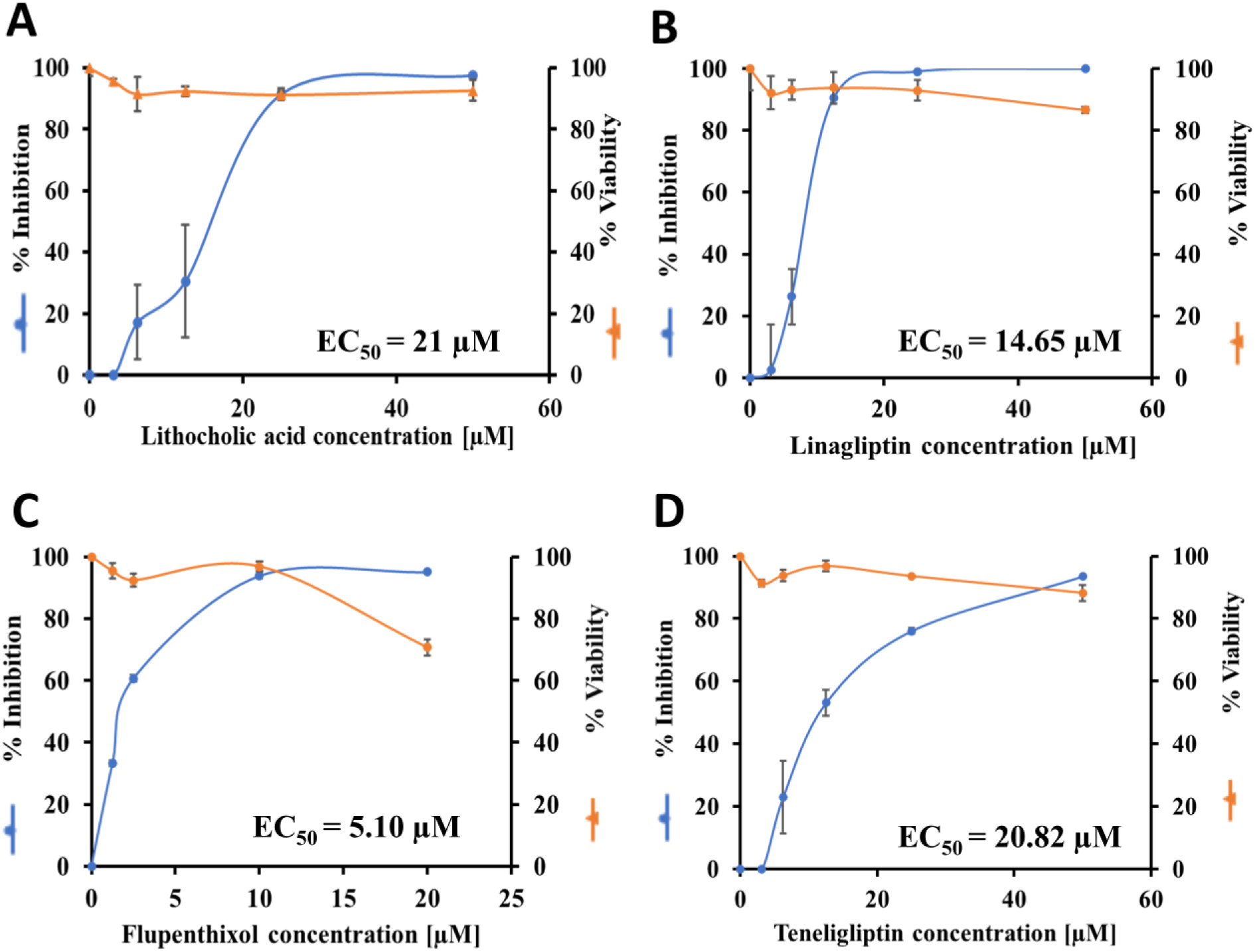
Antiviral efficacy of compounds against SARS-CoV-2 infection in Vero cells. (A-D) Vero cells were treated with increasing concentration of compounds and then infected with SARS-CoV-2 at a multiplicity of infection (MOI) of 0.01. Cell culture supernatant was harvested and the virus replication inhibition was quantified by qRT-PCR at 48 hpi. EC_50_ values for virus inhibition were estimated from dose–response curves of percentage inhibition versus compound concentration. Representative data are shown from duplicate readings and the final graph was plotted for percentage inhibition of cells in the presence and absence of compounds (n = 2). Data represent mean ± SD.

### Virus replication inhibition in human cell lines

To ensure that the observed antiviral efficacies of selected compounds were not restricted to Vero cells, the compound efficacies were further evaluated on two additional human cell lines expressing ACE2, HEK293T-ACE2 and A549-ACE2 that support SARS-CoV-2 replication. Cytotoxicity assay was performed in both HEK293T-ACE2 and A549-ACE2 cells to determine the working concentrations of the compounds. Lithocholic acid was toxic in HEK293T-ACE2 cells at a concentration of ∼25 µM, and flupenthixol at ∼20 µM concentration led to ∼20% loss in cell viability in both cell lines (Figure S4). Based on this, the following highest non-toxic doses were chosen: 50 µM teneligliptin, 50 µM linagliptin, 25 µM lithocholic acid, and 10 µM flupenthixol. *In vitro* cell based antiviral assays were performed and the dose-titration analysis revealed that 3 out of 4 compounds inhibited replication of SARS-CoV-2 in one or both of these cell lines at potencies equivalent to or greater than those that was observed in case of Vero cells. Results showed consistent inhibition of vRNA loads by a scale of ∼ 3 log_10_, in the presence of 50 µM linagliptin in HEK293T-ACE2 (EC_50_= 7.4 µM) and A549-ACE2 (EC_50_= 0.24 µM) cells respectively (Figure 4A, B, D, E). Lithocholic acid did not show any antiviral effects in HEK293T-ACE2 cells, but inhibited viral load by ∼1.5 log_10_ (EC_50_ = 0.6 µM) in A549 ACE2 cells (Figure 4A, C, D, F). Flupenthixol inhibited viral load by ∼ 3 log_10_ in HEK293T-ACE2 cell line but failed to show any effect in A549-ACE2 cells (Figure 4A, D). A possible reason for the differential efficacy of some compounds in different cell lines could be species-specific or cell-type-specific mechanism of action of a given compound or by differences in drug metabolism (Biering et al., 2021; Riva et al., 2020).

**Figure 4.**
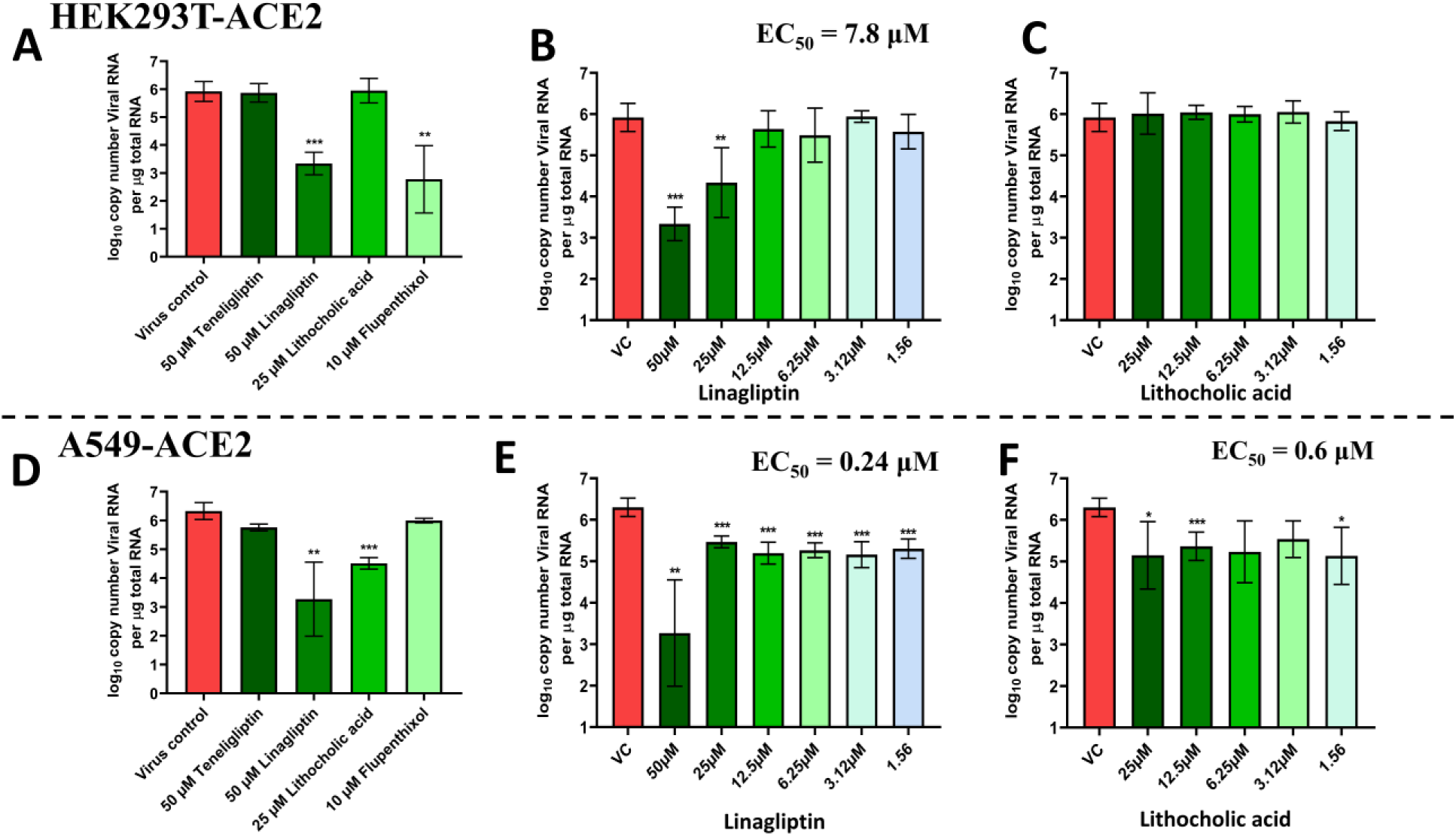
Antiviral efficacy of compounds against SARS-CoV-2 across multiple cell lines. (A-F) Cells were pre-treated with indicated drug concentrations for 3 h, washed and infected with 0.01 MOI SARS-CoV-2 for 2 h in the absence of drugs. After further washing, cells were infected with infection media containing appropriate inhibitor concentrations and after 48 h viral RNA load was estimated from total cellular RNA by qRT PCR. (A) shows results for HEK293T-ACE2 cells treated with single non-toxic concentrations of teneligliptin, linagliptin, lithocholic acid or flupenthixol, (B, C) show data for increasing concentrations of linagliptin and lithocholic acid respectively. (D-F) represent similar datasets in A549-ACE2 cells. Results represent data from 2 independent experiments. *p < 0.05, **p < 0.01, ***p < 0.001 using Brown-Forsythe and Welch ANOVA with Dunnett’s T3 multiple comparison tests. Error bars represent mean ± SD.

### Effects of compounds on cytokine expression and IFN responses

Infection by SARS-CoV-2 induces a hyper-inflammatory response driven by the uncontrolled release of TNF-α, IL-6, IL-1, IL-8, CXCL10 or delayed type I interferon response (Costela-Ruiz et al., 2020; Hu et al., 2021; Lan et al., 2021). Therefore, to understand the effects of identified compounds on the mRNA expression levels of cytokines, qRT-PCR was employed to quantify cytokine expression levels in treated cells. Interestingly, lithocholic acid and linagliptin were observed to inhibit the expression of IL6 (to a greater extent) and TNFα in HEK293T cells stimulated by polyinosinic:polycytidylic acid (poly I:C), a synthetic analog of dsRNA that mimics virus infection and supports that treatment with lithocholic acid or linagliptin could have an effect on virus induced inflammatory responses (Figure 5A and B). The PLpro of SARS-CoV-2 antagonizes the type-1 IFN innate immune responses of the host cell by reducing the amounts of ISGylated IRF3 or by deubiquitination of stimulator of interferon genes (STING) (Cao et al., 2023; Chen et al., 2014; Yuan et al., 2022). Consistent with these findings and as expected, the results of this assay also revealed that PLpro of SARS-CoV-2 suppressed the levels of IFNβ (Figure 5C)(Crudele et al., 2022; Shin et al., 2020). Further experimental analysis revealed that the compounds lithocholic acid and flupenthixol were able to restore the levels of interferon-beta (IFNβ) in HEK293T cells transfected with SARS-CoV-2-PLpro (Figure 5C). By contrast, no effect was observed on the expression levels of IFNβ when cells were treated with linagliptin and teneligliptin (Figure 5C). To investigate that the restored levels of IFNβ is due to inhibition of PLpro, an additional experiment was performed in which a luciferase reporter assay was used to test the effects of drugs on type-1 interferon induction using an IFNβ promoter-driven firefly luciferase reporter plasmid (IFN-Beta_pGL3) and a control plasmid which constitutively expresses renilla luciferase gene (pRL-TK). RIG-I 2CARD was used as an inducer of type-1 IFN induction at a very early step of the signaling pathway upstream of nuclear translocation of IRF3 and expression of interferon stimulated genes (ISGs). The plasmids were co-transfected with plasmid expressing SARS-CoV-2 PLpro to test the effect of the drugs to rescue the viral protein induced inhibition of IFN-β production. Having demonstrated a role for expression of SARS-CoV-2 PLpro in attenuating host antiviral IFN pathways, it was anticipated that inhibition of PLpro by identified compounds would reverse this process. Interestingly, treatment with lithocholic acid caused ∼4-fold increase of IFN-β production signifying that the compound was able to restore the suppressed type 1 interferon responses and these drugs could not rescue the PLpro induced IFN inhibition (Figure 5D and E).

**Figure 5.**
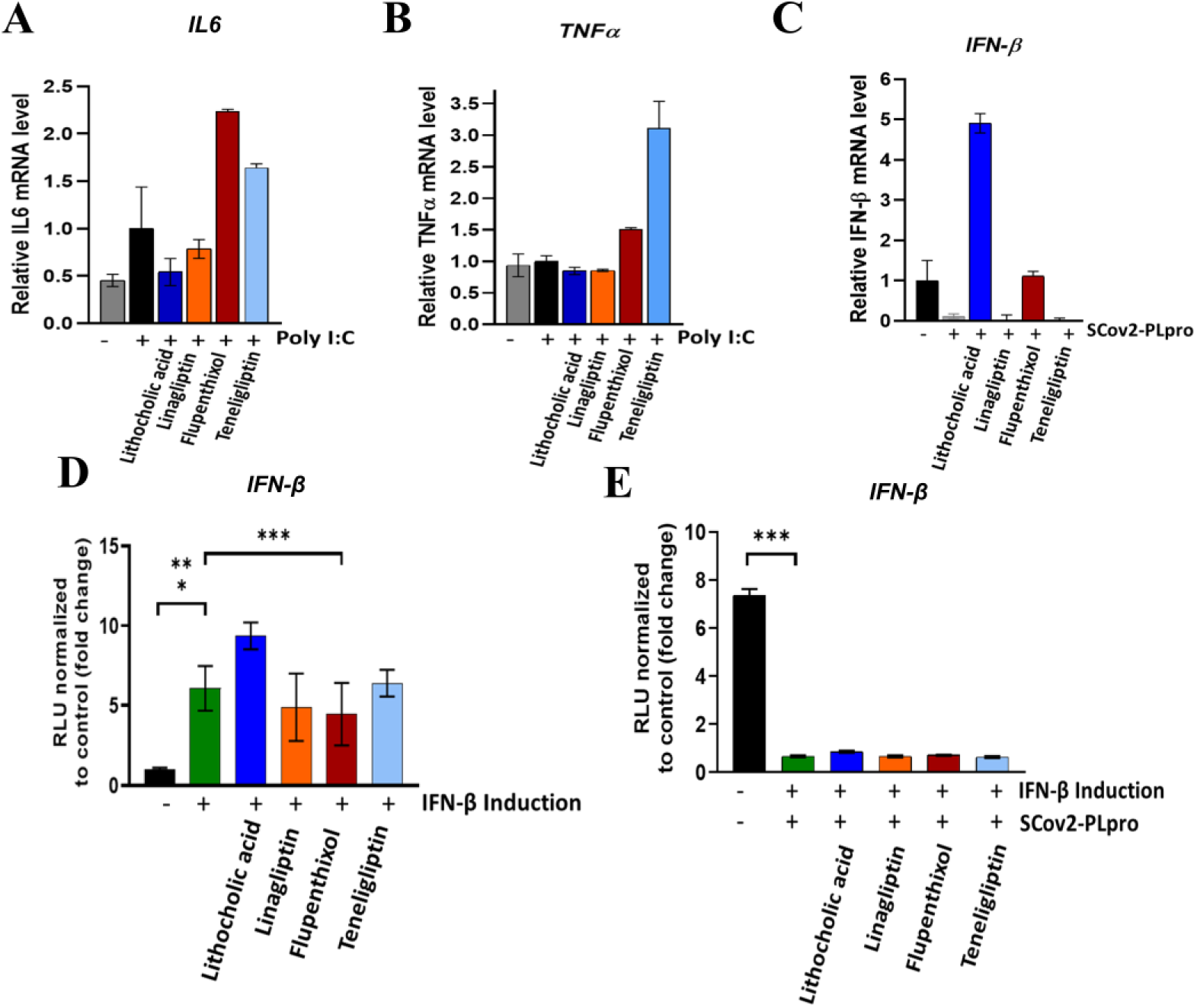
Effects of compound treatment on inflammatory cytokines and delayed type I interferon response. (A, B) HEK293T cells were stimulated with Poly I:C for 6 h and were treated with 30 µM of lithocholic acid (blue), 30 µM of linagliptin (orange), 10 µM of flupenthixol (red), and 30 µM of teneligliptin (light blue). (C) For evaluation of IFN-β levels, the cells were transfected with plasmid encoding SARS-CoV-2 PLpro followed by stimulation with poly I:C and treatment with compounds. RNA was isolated after 24 h and gene expression analysis for cytokines was performed using qRT-PCR. (D, E) HEK292T cells were transfected with IFNβ-luc firefly luciferase reporter plasmid and pRL-TK Renilla luciferase reporter plasmid, along with plasmid expressing SARS-CoV-2 PLpro or empty vector. RIGI-2 CARD was used as an inducer of type-1 interferon production. 3 h post transfection, cells were treated with different drugs 25 µM of lithocholic acid, 50 µM of linagliptin, 10 µM of flupenthixol, and 50 µM of teneligliptin, and 24 h later, cells were collected, and luciferase expression was analyzed by dual luciferase assay. (D) shows results for the effects of drugs on IFN-β promoter activity. (E) shows effects of drugs in rescuing SARS-CoV-2 PLpro induced inhibition of IFN-β production. Results represent data from 2 independent experiments. ***p < 0.001; using One-way ANOVA with Dunnett’s T3 multiple comparison tests. Error bars represent mean ± SD.

### Crystal structures of PLpro-inhibitor complexes

For elucidation of inhibitory mechanism of identified inhibitors, we made efforts to determine the crystal structures of PLpro in complex with inhibitors:lithocholic acid, linagliptin, teneligliptin, and flupenthixol. The data was obtained for the complex structures of PLpro:linagliptin and PLpro:lithocholic acid, which were solved in P3_2_21 space group at a resolution of 2.7 Å and 2.3 Å, respectively with one biologically active monomer in the asymmetric unit. Similar to previously reported apo structures of SARS-CoV-2 PLpro, the PLpro is structurally characterized as a monomer consisting of N-terminal ubiquitin-like domain (UBL) formed by β strands (β1– β3), the α-helical (α2–α7) thumb domain, the β-stranded finger domain, spanning β4– β7, and the palm domain (β8–13) (Figure 1B, 6A) (Fu et al., 2021; Klemm et al., 2020; Shen et al., 2022; Swaim et al., 2021). Located at the interface of the thumb–palm sub-domains lie a canonical catalytic triad active site made up of Cys111, His272, and Asp286 (Figure 1C) (Fu et al., 2021; Klemm et al., 2020; McClain and Vabret, 2020; Swaim et al., 2021). The fingers subdomain is structurally the most intricate domain, featuring a zinc-binding site where a structural zinc-ion is coordinated by four cysteines (Cys189, 192, 224, and 226) (Figure 1B, 6A) (Fu et al., 2021; Gao et al., 2021; Osipiuk et al., 2021). An important mobile β-turn/loop (BL2), positioned between β11–12 strands (residues 267-271), is a highly a flexible loop adjacent to the active site that recognizes and binds to the P2–P4 residues of the LXGG motif of the substrate (Gao et al., 2021; Osipiuk et al., 2021).

### Structural insights into PLpro-linagliptin complex

To elucidate the inhibitory mechanism of PLpro by linagliptin, we comprehensively analyzed the crystal structure of PLpro in complex with linagliptin. In the solved complex structure, linagliptin is found to accommodate the S2-binding site of PLpro, near the thumb domain (Figure 6A). In-depth structure analysis revealed that linagliptin interacts with the side chains of residues: Phe69, His73, Gln174, Ala176, Leu178, and Asp179 through hydrophobic interactions (Figure 6A-C). The side chains of Asn128, Gln174 and His175 are involved in H-bonding with the xanthine and purinedione skeleton of linagliptin (Figure 6A and B). Superimposition of PLpro structures complexed with linagliptin and ISG15 (PDB ID: 6YVA) revealed that linagliptin engages a region on PLpro that overlaps with Ub/ISG15 substrate binding site, and thus, is expected to block interaction of PLpro and ISG15/Ub substrates (Figure 6D and E). Interestingly, the 3-amino-piperidine ring and the purinedione skeleton of linagliptin bind to Phe69, His73, and Asn128 within the S2 Ub/ISG15-binding pocket of PLpro and these are the key residues that mediate hydrophobic interactions with ISG15 and ubiquitin substrates (Figure 6B-E) (Békés et al., 2016; Klemm et al., 2020; Wydorski et al., 2023). The critical residues Ser22, Met23, and Glu27 available on the binding surface of ISG15 are no longer able to access the S2 pocket of PLpro as the bound linagliptin molecule blocked the site (Figure 6E). Consequently, bound linagliptin is creating steric hindrance and blocked the access of the N-terminal domain of the substrates ISG15/Ub.

**Figure 6.**
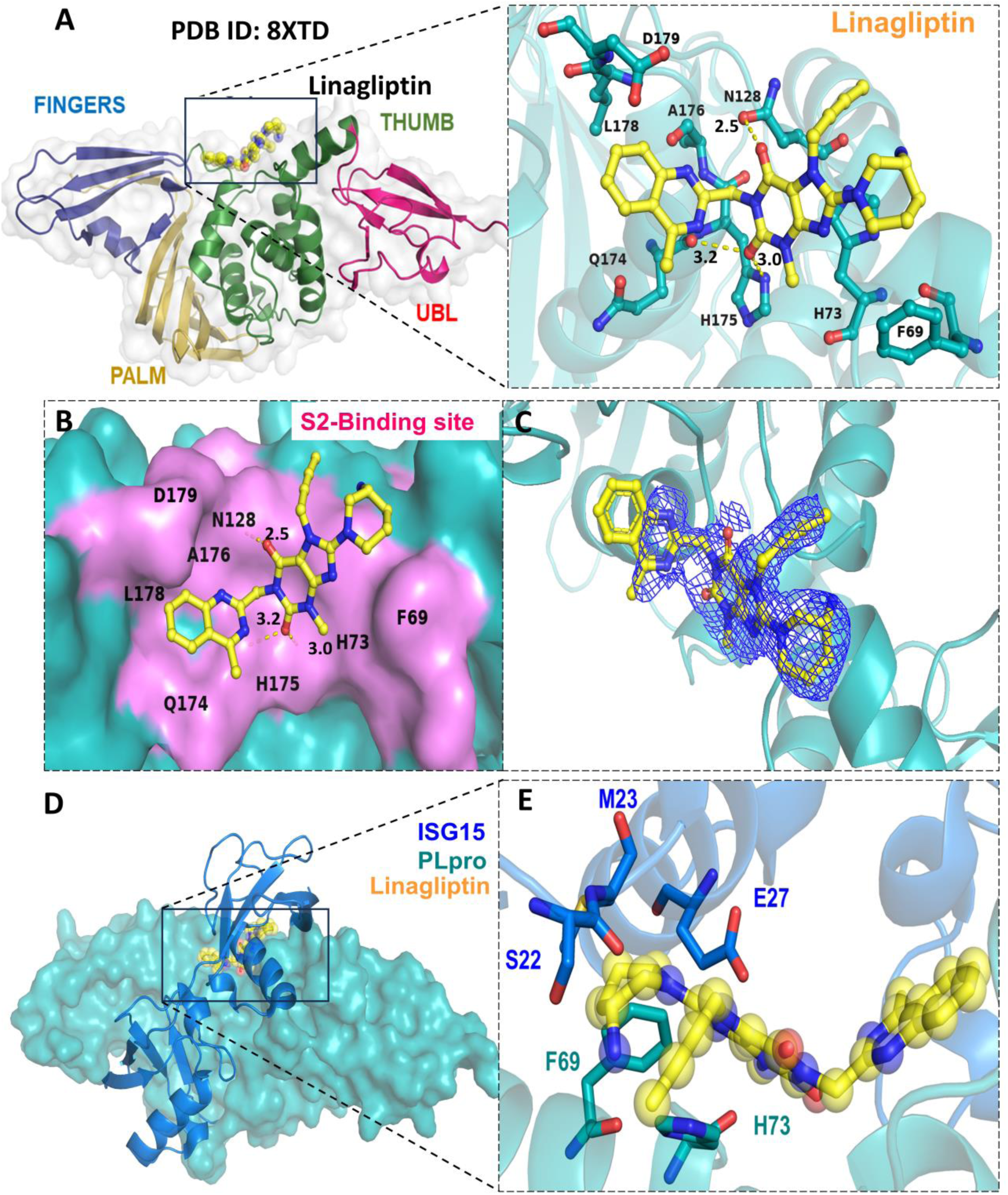
(A) Crystal structure of SARS-CoV-2 PLpro in complex with identified compound linagliptin (PDB ID: 8XTD) with PLpro shown as cartoon representation and the linagliptin shown as yellow spheres. The identified compound linagliptin binds at the Ub/ISG15 S2-binding pocket. The thumb domain (residues 61-180) is colored as green (residues 61-180), the palm (residues 239-315) as yellow, the finger (residues 181-238), and the UBL domain (residues 1-60) as red. (B) An enlarged view depicting intermolecular interactions of linagliptin to the S2-binding pocket of PLpro. The compound is shown as yellow sticks and the PLpro residues involved in interactions are depicted as teal green sticks. Dashed yellow lines show hydrogen bonds. (C) The electron density Fo-Fc map (contour level = 3 σ) is shown as blue mesh. (D, E) Superimposition of the crystal structure of SARS-CoV-2 PLpro:linagliptin (PDB ID: 8XTD) complex with crystal structure of SARS-CoV-2 PLpro in complex with ISG15 (PDB ID: 6YVA, ISG15 in blue cartoon). SARS-CoV-2 PLpro is depicted as teal green and the identified compound linagliptin is represented as yellow spheres. All structure analysis and figure preparation were carried out with PyMOL.

A difference Fo-Fc electron density map contoured at 3σ showed clear electron density (Fig 6C) and allowed unambiguous tracing of bound linagliptin molecule at the S2 site. However, the electron density of the terminal ring of quinazoline moiety was partial despite multiple attempts. This prompted us to evaluate the stability and intactness of linagliptin molecule in the condition used for performing soaking experiments along with the effects of crystallization buffers on integrity of the compound. HPLC analysis was performed using samples of linagliptin incubated in crystallization buffer at different time points and a single elution peak was obtained at retention time ∼3.2 min for the incubated samples and the UV spectra gave peak at ∼230 nm for samples. The obtained results were same as that of the standard solution of linagliptin suggesting that linagliptin does not get degraded in the crystallization buffers and the compound is structurally intact (Figure S6). In conclusion, a slightly weak electron density of this aromatic ring of quinazoline moiety could presumably be due to a high flexibility of this ring (Davis et al., 2003).

### Structural insights into PLpro-lithocholic acid complex

For understanding the molecular basis of SARS-CoV-2 PLpro inhibition by lithocholic acid, the three-dimensional structure of PLpro bound to lithocholic acid was determined (PDB ID: 8X1X). The electron densities in the structure revealed that three molecules of lithocholic acid occupy three binding sites in SARS-CoV-2 PLpro, which included the substrate binding-cleft near the BL2 groove and the Ub/ISG15 S2-binding pocket (Figure 7). At the S2-binding pocket, lithocholic acid I form hydrophobic interactions with the side chains of Phe69, His73, Asn128, Gln174, His175, Ala176, Asn177, Leu178, and Asp179 (Figure 7A-C). Interaction of lithocholic acid I with the substrate interacting residues of PLpro (Phe69, His73, Asn128) reveals molecular basis of inhibition mechanism of PLpro by lithocholic acid I, disrupting the protein-protein interaction (PPI) between PLpro and ISG15 or di-Ub (Figure 7B, C). Superimposition of PLpro:lithocholic acid I complex structure over SARS-CoV-2 PLpro:ISG15 structure suggests disruption of interactions between PLpro and ISG15. Remarkably, in the presence of lithocholic acid I, the essential residues of ISG15, Ser22, Met23, Ser26, and Glu27, which are known to interact with the S2 pocket of PLpro, are no longer accessible to form contacts with PLpro (Figure 7B). To our surprise, a second binding site for lithocholic acid II was observed at a novel site near the zinc-finger motif (Figure 7D). The interactions of lithocholic acid II are stabilized by hydrophobic interactions with Tyr213, Lys217, Thr257, Tyr305, Lys306, Glu307, Asn308, and Ser309 (Figure 7D). Additionally, lithocholic acid II forms H-bond interaction with Glu214 further stabilizing the complex (Figure 7D).

**Figure 7.**
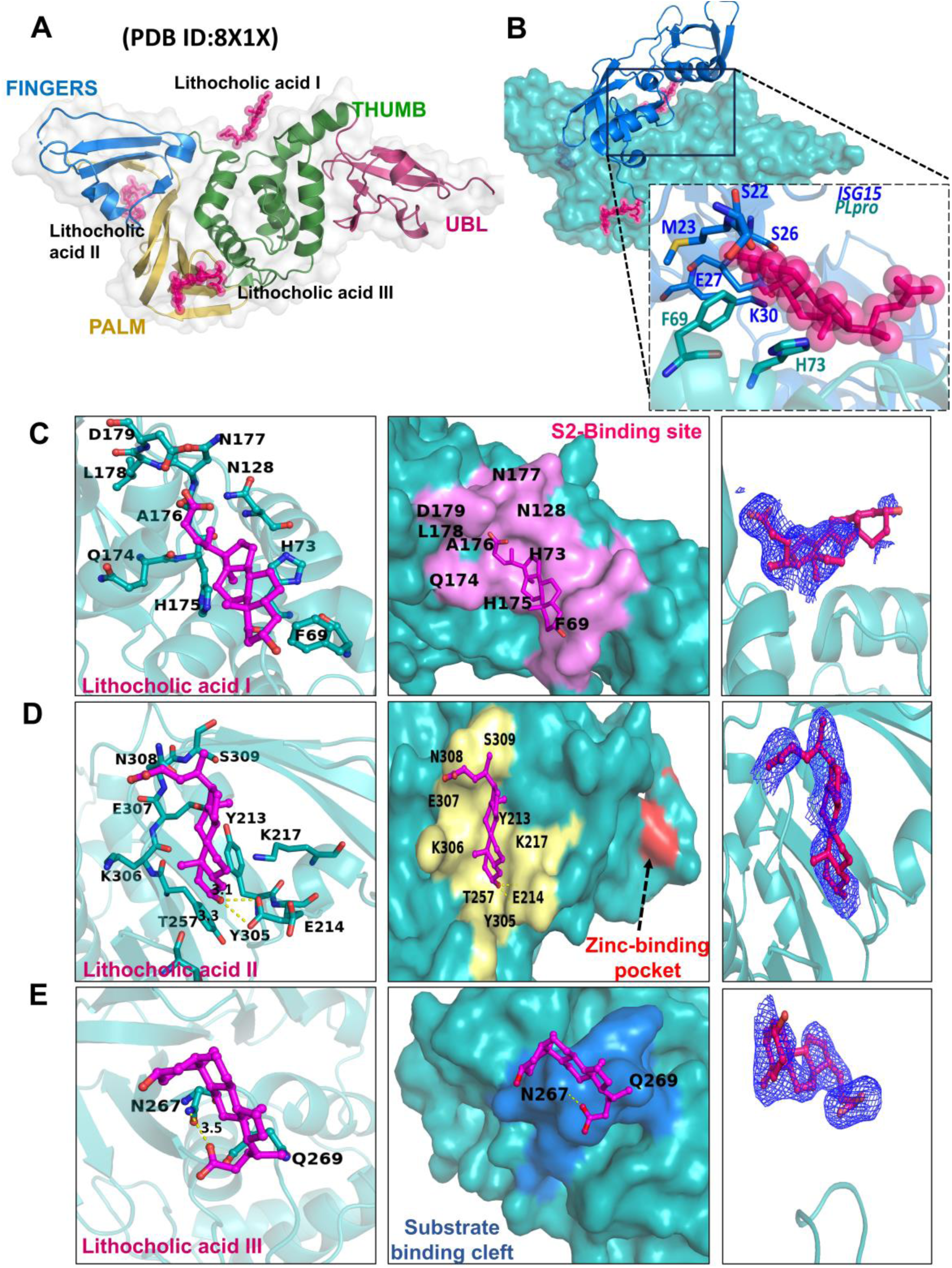
(A) Crystal structure of SARS-CoV-2 PLpro in complex with lithocholic acid where PLpro is depicted as cartoon representation and the lithocholic acid is shown as pink spheres. The identified compound binds at three sites: lithocholic acid I (Ub/ISG15 S2-binding site), II (a novel site near the zinc-binding pocket), and III (substrate binding cleft). The thumb domain is colored as green (residues 61-180), the palm domain as yellow (residues 239-315), the finger domain as blue (residues 181-238), and the UBL domain (residues 1-60) as red. (B) Superimposition of the crystal structure of SARS-CoV-2 PLpro:lithocholic acid complex with crystal structure of SARS-CoV-2 PLpro in complex with mouse-ISG15 (PDB ID: 6YVA, ISG15 in blue cartoon). SARS-CoV-2 PLpro is depicted as teal green surface and the identified compound lithocholic acid is represented as pink spheres at three sites. An enlarged view depicting intermolecular interactions of (C) lithocholic acid I to the S2-binding pocket, (D) lithocholic acid II to a novel site near the zinc-binding pocket, (E) and lithocholic acid III to substrate-binding pocket. The compound is shown as pink sticks and the PLpro residues involved in binding are depicted as teal green sticks. Dashed yellow lines show hydrogen bonds. The electron density Fo-Fc map (contour level = 3 σ) is shown as blue mesh. All structure analysis and figure preparation were carried out with PyMOL.

In addition, a third molecule of lithocholic acid III, occupies the substrate-binding cleft pocket forming interactions with Asn267 and Gln269, the two prime residues of the flexible BL2 loop (Figure 7E). Conformational changes in the BL2 loop are reported to regulate the binding, substrate accessibility, and enzymatic activity of PLpro (Fu et al., 2021; Napolitano et al., 2022; Osipiuk et al., 2021). Asn267 and Gln269 are reported to accommodate the C-terminal leucine residue at P4 position of PLpro substrates. Interestingly, analysis of the BL2 loop suggests that upon binding with lithocholic acid III, the main chain atom of Asn267 forms H-bond with inhibitor (Figure 7E). The ligand is presumed to stabilize the inhibited state of PLpro after establishing relevant interactions with Gln269 (BL2-loop) and Asn267 (substrate binding site). A similar set of interactions were previously observed in other reported crystal structures of PLpro (PDB ID: 7LBR, 7JIR, and 7JIW) and specific interaction with Asn267 and Gln269 was shown to be essential for inhibition. The binding of lithocholic acid does not induce significant conformational changes in the BL2 when compared with unliganded apo form of PLpro (PDB ID: 6WZU). The most dramatic change was observed at the residue Gln269. In the apo structure, the side chain of Gln269 swings away from the pocket (Figure 7E). In the lithocholic acid-bound structure, the side chain of Gln269 rotates inwards towards the substrate-recognition cleft, thus narrowing the substrate-binding cleft and clamping the inhibitor to the protease (Figure 7E). The crystallographic data and refinement statistics are listed in Table 2.

**Table 2.**
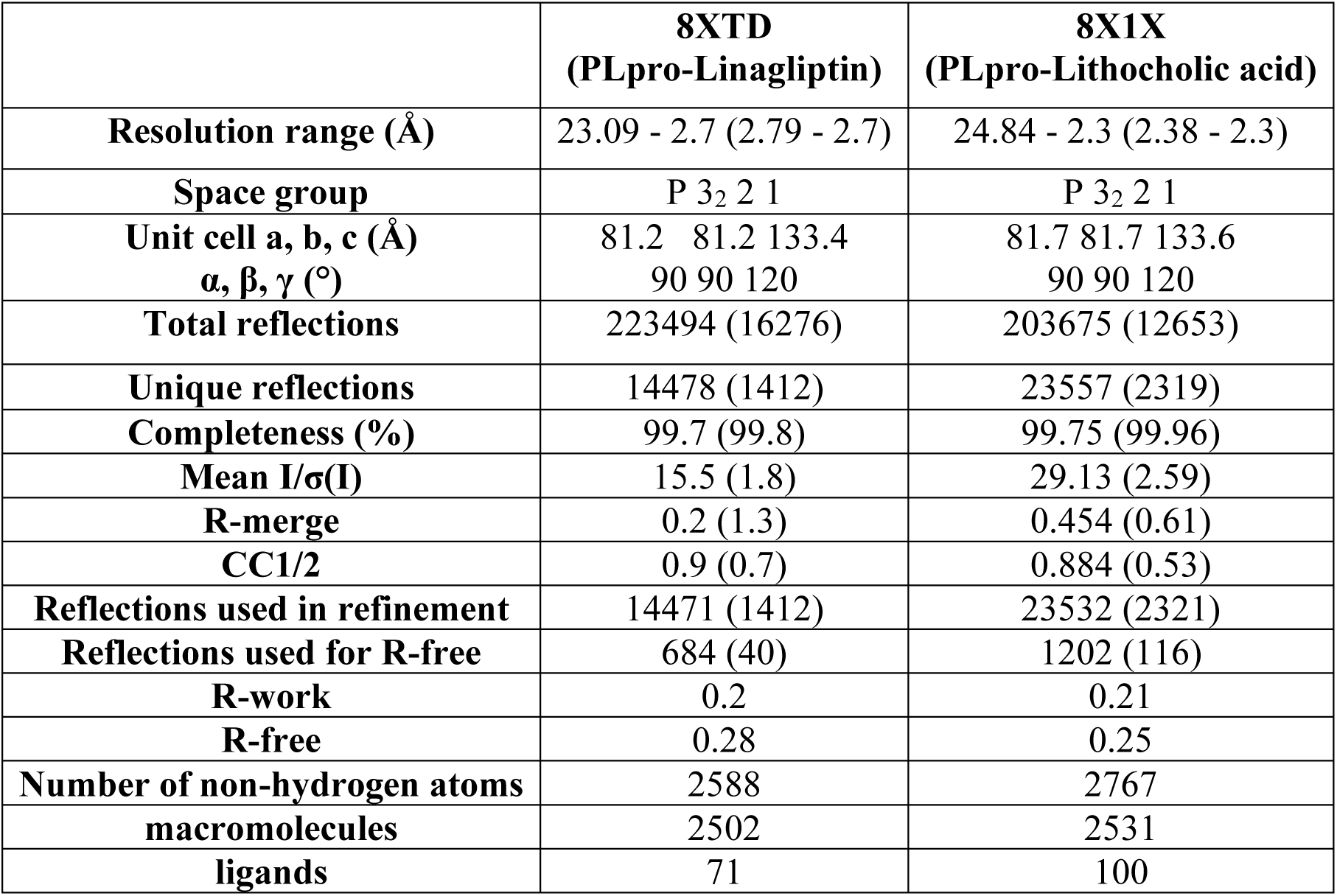
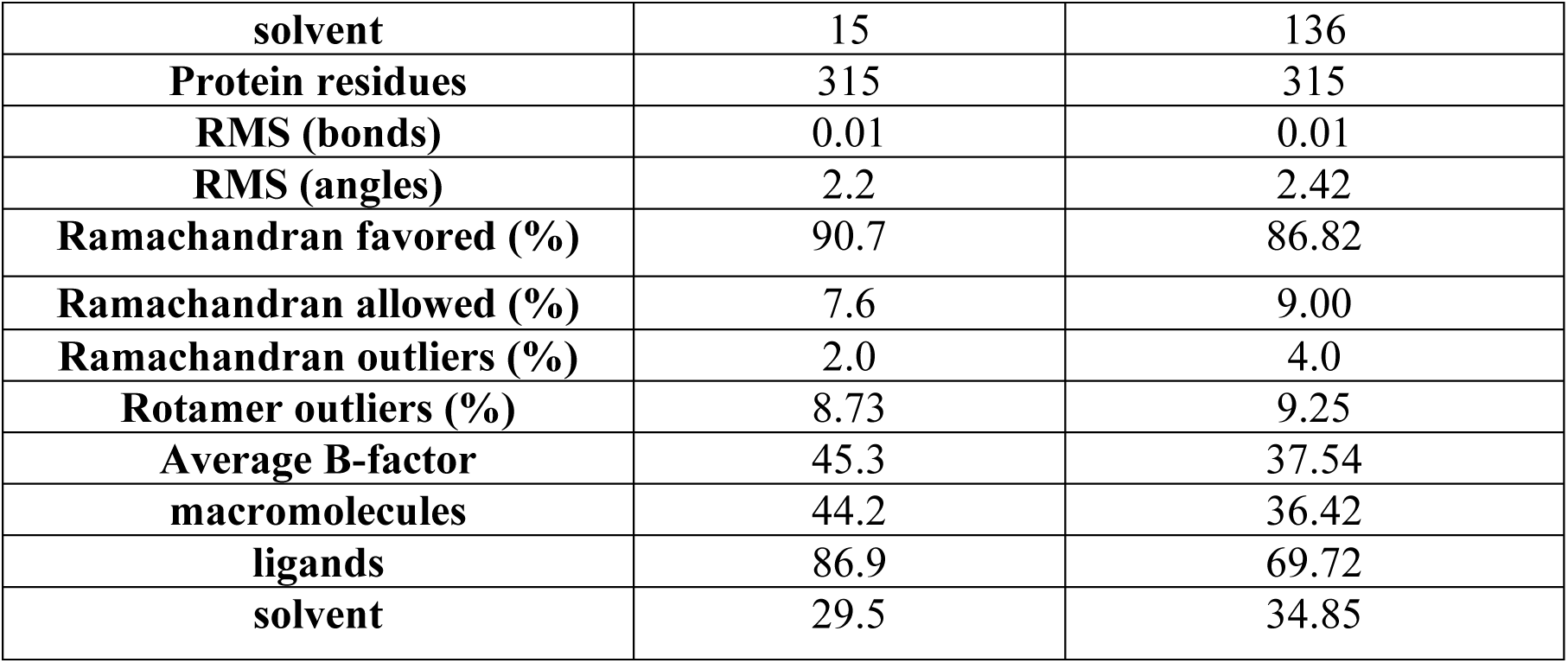
Data collection and refinement statistics.

|  | **8XTD**<br>**(PLpro-Linagliptin)** | **8X1X**<br>**(PLpro-Lithocholic acid)** |
| --- | --- | --- |
| **Resolution range (Å)** | 23.09 - 2.7 (2.79 - 2.7) | 24.84 - 2.3 (2.38 - 2.3) |
| **Space group** | P 3<sub>2</sub> 2 1 | P 3<sub>2</sub> 2 1 |
| **Unit cell a, b, c (Å)**<br>**$\alpha$, $\beta$, $\gamma$ (°)** | 81.2 81.2 133.4<br>90 90 120 | 81.7 81.7 133.6<br>90 90 120 |
| **Total reflections** | 223494 (16276) | 203675 (12653) |
| **Unique reflections** | 14478 (1412) | 23557 (2319) |
| **Completeness (%)** | 99.7 (99.8) | 99.75 (99.96) |
| **Mean I/$\sigma$(I)** | 15.5 (1.8) | 29.13 (2.59) |
| **R-merge** | 0.2 (1.3) | 0.454 (0.61) |
| **CC1/2** | 0.9 (0.7) | 0.884 (0.53) |
| **Reflections used in refinement** | 14471 (1412) | 23532 (2321) |
| **Reflections used for R-free** | 684 (40) | 1202 (116) |
| **R-work** | 0.2 | 0.21 |
| **R-free** | 0.28 | 0.25 |
| **Number of non-hydrogen atoms** | 2588 | 2767 |
| **macromolecules** | 2502 | 2531 |
| **ligands** | 71 | 100 |
| **solvent** | 15 | 136 |
| **Protein residues** | 315 | 315 |
| **RMS (bonds)** | 0.01 | 0.01 |
| **RMS (angles)** | 2.2 | 2.42 |
| **Ramachandran favored (%)** | 90.7 | 86.82 |
| **Ramachandran allowed (%)** | 7.6 | 9.00 |
| **Ramachandran outliers (%)** | 2.0 | 4.0 |
| **Rotamer outliers (%)** | 8.73 | 9.25 |
| **Average B-factor** | 45.3 | 37.54 |
| **macromolecules** | 44.2 | 36.42 |
| **ligands** | 86.9 | 69.72 |
| **solvent** | 29.5 | 34.85 |

### Linagliptin exhibits *in vivo* antiviral efficacy in mouse model of SARS-CoV-2 infection

Linagliptin, the lead compound was the most promising based on both the biophysical binding assessments and *in vitro* anti-SARS-CoV-2 studies. Therefore, the *in vivo* antiviral efficacy of the compound was investigated using a mouse-adapted SARS-CoV-2 model (MA10). To this end, mice were intranasally infected with 1×10^5^ PFU of mouse-adapted SARS-CoV-2 (MA10) 6 h before the first dose. Challenged mice received treatments with linagliptin twice daily for 4 days via intraperitoneal (i.p.) and oral (p.o.) routes at doses of 30 mg/kg and 60 mg/kg. Molnupiravir was used as a positive control and dosed orally and intraperitoneally at 100 mg/kg and 200 mg/kg. SARS-CoV-2 lung titers were quantified for the treatment group and were compared to vehicle and molnupiravir control (Figure 8B). Overall, both i.p and oral administration of linagliptin (30 mg/kg and 60 mg/kg) displayed promising antiviral efficacy of linagliptin against SARS-CoV-2 with more effective outcomes in oral administration than the intraperitoneal route (Figure 8B and C). The antiviral effect of linagliptin was evident with over a log viral titer reduction in lung viral load (60 mg/kg i.p and 30 mg/kg oral) relative to the control infected groups (Figure 8B). Histopathological analysis revealed that oral administration of linagliptin (30 mg/kg and 60 mg/kg) significantly reduced the severity of lung lesions, including peribronchiolar and perivascular infiltration, bronchiolar epithelial desquamation, and alveolar septal infiltration, compared to the control infected groups (Figure 8C and D).

**Figure 8.**
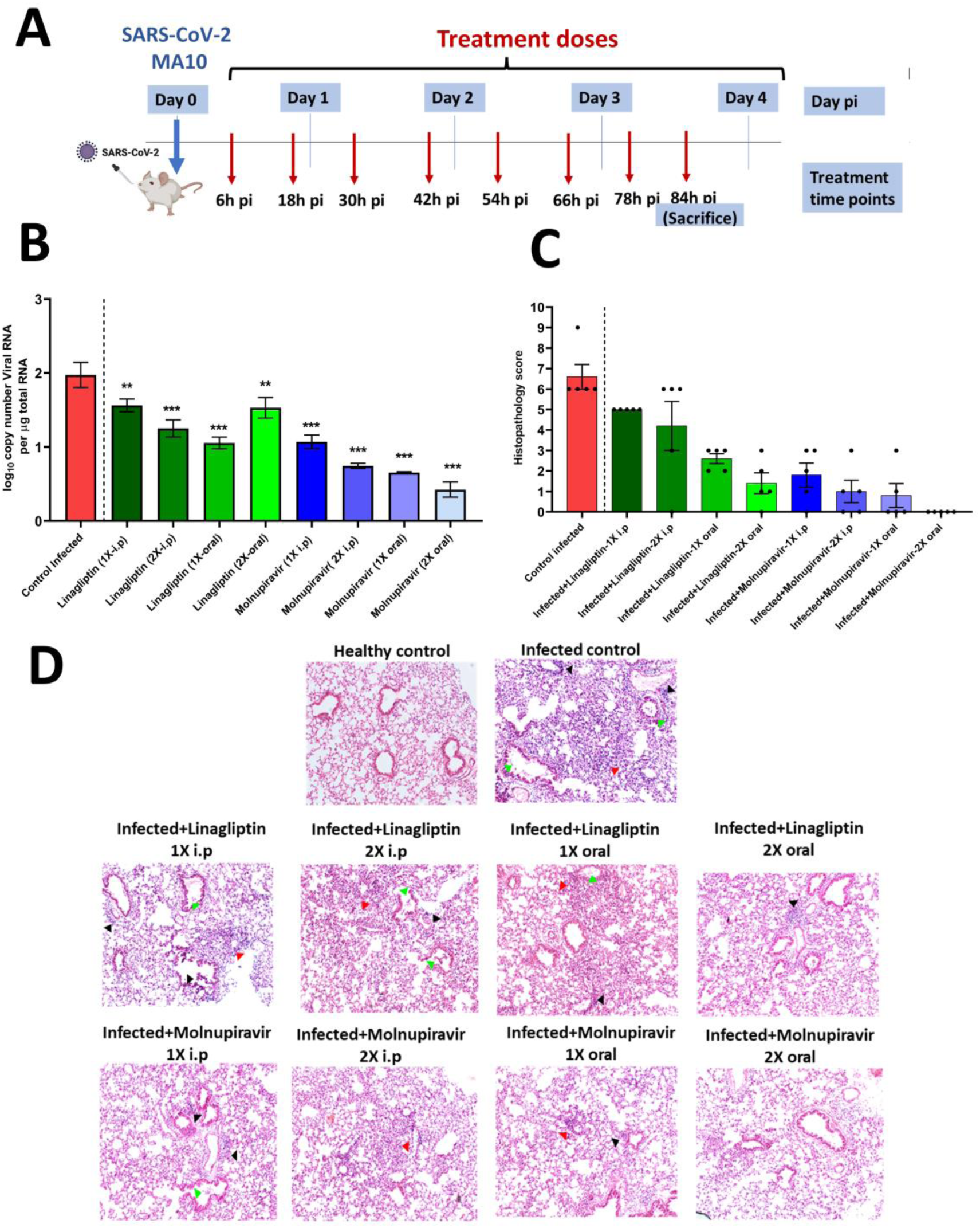
Linagliptin shows in vivo antiviral efficacy in mouse-adapted model of SARS-CoV-2 (MA-10). (A) A schematic view of the experimental plan. Female mice were dosed with two different concentrations of linagliptin and molnupiravir starting from 6 h post infection until the day of sacrifice. (B-C) On day 4 post infection, lungs were harvested for estimating viral RNA load and histology analysis. (D) Histopathological scoring of the lung tissue. Lung histopathology lesions (10X) showing peribronchiolar and perivascular infiltration (black arrow head), bronchiolar epithelial desquamation (green arrow head) and alveolar septal thickening (red arrow head).

## Discussion

The rapid rollout of vaccines played a crucial role in battling the pandemic of COVID-19 (Li et al., 2022; Mello et al., 2022). However, the rise of emerging variants, breakthrough infections, and drug resistance underscores the critical need for orally bioavailable broad-spectrum antivirals as a part of pre-preparedness plan. Drug repurposing provides an alternative approach for quick identification of potential drugs to combat COVID-19, and targeting PLpro offers additional broad-spectrum advantages owing to its druggability and highly conserved substrate-binding pocket. Several host proteases, including USP2 and USP7 exhibit the same deconjugating activity and structural homology as host-encoded DUBs, marking their similarity as a noteworthy target to be explored (Figure 1). In addition to the protease activity, PLpro possesses DUB and deISGylating activities as an evasion mechanism against the host antiviral immune response. To evade host immunity, PLpro antagonizes ISG15 dependent activation of MDA5 and RIG1 protein through its DUB activity (Liu et al., 2021). A dysregulated interferon mediated antiviral response could lead to pro-inflammatory cytokine storm in the COVID-19 patients, a major cause of mortalities among infected patients (Liu et al., 2021). Thus, targeting PLpro might serve as a multifaceted approach that could avert SARS-CoV-2 infection either by inhibition of its proteolytic activity or by upregulation of innate immune response.

In search of a broad-spectrum antiviral drug, the present study investigated the antiviral efficacy of DUB inhibitors against PLpro, owing to the structural and functional similarities (Figure 1). Structure-based drug repurposing revealed five promising molecules targeting the Ub/ISG15 S2-binding pocket and the catalytic site of PLpro, from an *in-house* library of DUB inhibitors (Table 1, table S1, and Figure 1). The enzymatic assay and SPR binding kinetic studies against PLpro demonstrate that lithocholic acid, linagliptin, teneligliptin, and flupenthixol hold tremendous promise as a new class of antivirals against PLpro (Figure 2). Interestingly, ligand-binding experiments using SPR were in line with the enzymatic inhibition studies with observed K_D_ values in the micromolar range (lithocholic acid -PLpro: 79.3 μM; linagliptin-PLpro: 72.6 μM; flupenthixol-PLpro: 258.1 μM) (Figure 2). The selected compounds efficiently inhibited not only the proteolytic roles of PLpro but also the viral replication (Figure 2 and 3). Concentration-dependent reduction in viral copy number and infectious viral titer well below their cytotoxic concentration highlights the effective anti-SARS-CoV-2 activity of compounds in the initial experiments in Vero cells (Figure 3). Dose-titration data depicts that nearly all (3 out of 4) tested compounds significantly inhibited replication of SARS-CoV-2 in one or both HEK293T-ACE2 and A549-ACE2 cell lines at potencies equivalent to or greater than those that was observed in case of Vero cells (Figure 4). Though teneligliptin failed to produce discernible kinetic profiles in the SPR assay, its antiviral data suggested significant inhibitory effect against SARS-CoV-2, proposing a hint for other possible off-targets that could be explored (Figure 2 and 3). Linagliptin demonstrated the strongest inhibition in both A549-ACE2 and HEK293T-ACE2 cells with EC_50_ value of 0.24 and 7.4 µM, respectively (Figure 4). The cytotoxicity profile of these compounds in Vero cells, HEK293T-ACE2, and A549-ACE2 showed a CC_50_ value of > 50 μM marking them safe for use (Figure S4).

To obtain further mechanistic understanding of the mode of inhibition, structures of linagliptin and lithocholic acid in complex with PLpro were determined using X-ray crystallography (Figure 6 and 7). The data indicates that both linagliptin and lithocholic acid occupied the Ub/ISG15 S2-binding pocket of PLpro at the thumb domain, located at the interface between PLpro and ISG15 (Figure 6 and 7). Once linagliptin or lithocholic acid occupies the S2 interface between PLpro and ISG15 substrate, it is likely to disrupt the interaction between PLpro and natural substrates such as ISG15 or di-Ub. Furthermore, by blocking the access of Ub/ISG15 to the S2-binding pocket on PLpro, linagliptin or lithocholic acid offers potential to downregulate the deubiquitinase or deISGylase activity, thereby providing a novel strategy for structure-based inhibition of SARS-CoV-2 and its emerging variants (Figure 6 and 7). Of all the compounds tested, linagliptin exhibited significant inhibitory effects in biophysical, biochemical, and virus inhibition across multiple cell lines, and was thus selected as prime candidates for *in vivo* investigation in intranasally infected mouse-adapted MA10 SARS-CoV-2 model (Figure 8). Twice-daily oral and intraperitoneal administration of linagliptin (30 mg/kg and 60 mg/kg of body weight) to mice for 4-day period significantly reduced the lung viral RNA load, reduced weight loss, and ameliorated SARS-CoV-2 induced histopathological damages *in vivo* (Figure 8).

Previous studies on lithocholic acid report it to be a potent inhibitor of intestinal inflammatory responses (Ward et al., 2017). In another study, lithocholic acid is reported to exert antiviral action on replication of porcine delta coronavirus replication (PDCoV) (Kong et al., 2021). Recent clinical and experimental data on COVID-19 provides evidence for reduced mortality in infected patients with or without type 2 diabetes, after the use of gliptins (e.g., sitagliptin, alogliptin, etc.). Gliptins, a class of DPP4 inhibitors, antagonize SARS-CoV-2 virulence either by enhancing GLP-1 anti-inflammatory activity or by reducing overproduction of cytokines and downregulating the activity of macrophages (Strollo and Pozzilli, 2020)(Solerte et al., 2020). These classes of inhibitors may halt the disease progression to a hyperinflammatory state after viral infection.

In conclusion, the compounds identified in this study offer new perspectives for treatment of COVID-19. Given that the identified PLpro inhibitors acts through a novel mechanism, follow up characterization of these inhibitors in combination with Mpro and RdRp inhibitors may help to further mitigate the risk of drug-resistant viral evolution and enhance pandemic preparedness. Overall, linagliptin is a strong therapeutic candidate for further development as an oral anti-SARSCoV-2 antiviral.

## Experimental procedures

### Cells, viruses, and compounds

Vero and HEK293T cell line used in this study were procured from NCCS (Pune, India). The cells were maintained in high glucose Dulbecco’s-modified essential media (DMEM; HiMedia, India) augmented with 10% heat inactivated fetal bovine serum (FBS; Gibco, USA), 100 units of penicillin, and 100 mg streptomycin/mL. HEK293T cells expressing human ACE2 (HEK293T-ACE2) (NR-52511, BEI Resources, NIAID, NIH) and A549 cells (NCCS, Pune, India) transduced to stably express human ACE2 receptor (A549 ACE2), were used for SARS-CoV-2 infection. HEK293T-ACE2, HEK-293T, and A549-ACE2 cells were cultured in complete media prepared using DMEM (12100-038, Gibco) supplemented with 10% HI-FBS (16140-071, Gibco), 100 U/mL Penicillin-Streptomycin (15140122, Gibco) and GlutaMAX™ (35050-061, Gibco). Assessment of cytotoxic effect and the inhibitory potential of identified compounds against SARS-CoV-2 replication was carried out in Vero cells, HEK293T-ACE2, and A549-ACE2. HEK-293T cells (NCCS, Pune) were used for pro-inflammatory cytokine quantification assay and interferon (IFN) induction assay.

The wild type SARSCoV-2/Human/IND/CAD1339/2020 strain was isolated from laboratory confirmed COVID-19 patients in India. After genetic characterization by whole genome sequencing (GenBank accession no: MZ203529), the virus was passaged in Vero cells and supernatant was collected and clarified by centrifugation before being aliquoted for storage at −80 °C. Virus titer was measured in Vero cells by TCID_50_ assay. SARS-CoV-2 isolate Hong Kong/VM20001061/2020 (BEI resources NR-52282), 2020 (hereby referred as SARS-CoV-2) was propagated and titrated in VeroE6 cells. Teneligliptin, linagliptin, lithocholic Acid, vildagliptin, and flupenthixol (Cayman, India) were dissolved in DMSO and the stock solutions were prepared. The working concentration of drugs prepared in cell culture media was centrifuged at 10,000 rpm for 10 min, and the supernatant was used for all the assays. All procedures related to SARSCoV-2/Human/IND/CAD1339/2020 virus culture and antiviral studies were handled in Biosafety level 3 facility at Indian Veterinary Research Institute (IVRI), Izatnagar, Bareilly, following standard WHO guidelines. Experiments with SARS-CoV-2 isolate Hong Kong/VM20001061/2020 were performed at the viral biosafety level-3 facility, Indian Institute of Science, Bengaluru, India (Murgolo et al., 2021).

### Plasmid constructs

DNA sequence encoding the PLpro of SARS-CoV-2 was codon optimized for bacterial expression (Choudhary et al., 2023). PCR amplification of the gene encoding PLpro was carried out using synthesized gene (Invitrogen, Thermo Fisher scientific) as the template. After restriction digestion, the gene was cloned into the *Nde1* and *Xho1* restriction sites of pET28c bacterial expression vector using 5’-CAGCCATATGGAAGTTCGTACCATTAAAGT TTTTAC-3’ forward primer and 5’-GGTGCTCGAGCTATTTGATGGTGGTG-3’ reverse primer. The resulting plasmid was transformed into competent DH5α cells. These transformed cells were grown on Luria–Bertani (LB) agar plates supplemented with 50 μg/mL kanamycin with incubation at 37 °C overnight. Several colonies were picked and grown in 10 mL LB broth containing 50 μg/mL of kanamycin, followed by plasmid isolation. Using the wild type PLpro plasmid as the template, site directed mutagenesis was performed to generate an active site mutant (C111S) by an overlapping PCR strategy, using 5’-AAATGGGCAGATAATAACTCGTATCTGGCAACCG-3’ forward primer and 5’-AGTGCGGTTGCCAGATACGAGTTATTATCT-3’ as reverse primer. For mammalian expression, gene encoding PLpro was cloned into pcDNA3 vector between *BamH1* and *Xho1* restriction sites using forward primer 5’-ATGAGGATCCATGGAAGTTCGTACCATTAAAGTTTTTACCA-3’ and reverse primer 5’-GGTGCTCGAGCTATTTGATGGTGGTG-3’. The plasmids IFN-Beta_pGL3 (Addgene #102597); pRLTK; RIGI 2CARD; plx 304 empty vector (kind gift from Adolfo García-Sastre, Icahn School of Medicine at Mount Sinai, New York) were used for interferon induction experiment. All the clones were verified by DNA sequencing.

### Structural comparison of PLpro and USP protein

Pairwise comparisons of secondary structure elements were performed with the web-based Secondary Structure Matching program (SSM), available within the Protein Data Bank (PDBeFold) (Krissinel and Henrick, 2004). For assessing structural similarities, crystal structures of SARS-CoV-2 PLpro (PDB ID: 6W9C), HAUSP/USP7 (PDB ID: 2F1Z), and USP2 (PDB ID: 5XU8) were downloaded from RCSB-PDB (Berman et al., 2000). The structures were fed into the PDBeFold-SSM portal and the hits from each pairwise comparison are ranked by their RMSD values, Q-score, and the number of aligned residues. The 3D alignment file was downloaded and analyzed in PyMOL (Schrodinger LLC, 2015) to observe conserved loops and residues. To gain more insights into the conserved residues at the substrate binding pockets (Ub/ISG15 binding site), multiple sequence alignment of SARS-CoV-2 PLpro (PDB ID: 6W9C), HAUSP/USP7 (PDB ID: 2F1Z), and USP2 (PDB ID: 5XU8) was done using Clustal Omega/ESPript 3.0 (Robert and Gouet, 2014).

### Structure-based identification of inhibitors

An *in-house* library of compounds containing previously reported DUBs inhibitors and cyanopyrrolidine ring compounds was prepared, and screened against PLpro to identify potential antivirals against SARS-CoV-2 (Table S1). The SDF files for three-dimensional structures (3D) of all compounds were retrieved from PubChem (https://pubchem.ncbi.nlm.nih.gov/) database. 3D-crystal structure of PLpro of SARS-COV-2 (PDB ID: 6W9C) was retrieved from RCSB-PDB. Using the automated function of AutoDock Vina (Trott and Olson, 2010), refinement of protein was done by addition of polar hydrogen atoms and the protein was saved in PDBQT format for further docking studies.

Before performing structure-based virtual screening, all the ligands were energy minimized by Universal Force Field (UFF) using Open Babel in PyRx 0.8 (Dallakyan and Olson, 2015; O’Boyle et al., 2011). SDF files of all ligands were converted into PDBQT format for virtual screening against PLpro using an in-built Open Babel and AutoDock Vina module of PyRx 0.8. The grid centers for docking search space were set as X= −34.1, Y= 20.3, and Z=33.8, and the dimensions (Å) for X, Y, and Z axis were set as 36.4, 27.8, and 31 respectively. The size of the grid box was set centering on the substrate binding site for PLpro with a default exhaustiveness value of 8 for all ligands. A total of 9 different poses were generated for each ligand. Visual examination of the docked complexes possessing the highest binding affinities was carried out using PyMOL for analyzing hydrogen and hydrophobic interactions. Top-ranked compounds displaying interactions similar to GRL0617 for targeted site of PLpro, possessing high binding affinities, and low RMSD values (< 2 Å) were sorted out. Of these hits, the compounds that were recently reported as SARS-CoV-2 inhibitors in other studies were excluded from further studies.

Molecular docking of the ligands with PLpro of SARS-CoV-2 was carried out at two target sites, the substrate binding cleft near the BL2 groove and the Ub/ISG15 binding site (S2 site), using AutoDock 4.2.6 (Trott and Olson, 2010). Crystal structure of PLpro of SARS-CoV-2 (PDB ID: 6W9C) was downloaded from PDB and the PDBQT file of this energy minimized structure was used. AutoDock and AutoGrid tools available with AutoDock4 were used to calculate grid maps for pre-calculation of interaction energies of all interacting atoms. The center points for substrate-binding cleft were set as X= −34.4, Y= 27.1, and Z= 33.9; grid box dimensions were 100 Å ×100 Å×100 Å. Grid box dimensions for re-docking studies at the S2 Ub/ISG15 binding site were 80 Å ×60 Å×80 Å and grid center points were set as X= −17.6, Y=40.9, and Z= 11.5. Lamarckian genetic algorithm (GA) with a combination of maximum grid-based energy evaluation method was used for docking studies. The program was run for a maximum number of 25 genetic algorithm runs. All other parameters were set as default. A previously reported PLpro inhibitor GRL0617 was used as a reference control (Fu et al., 2021). Out of all the possible poses generated, the pose showing maximum binding affinity, hydrogen, hydrophobic interactions, and lowest RMSD values were chosen for detailed visual analysis using LigPlot and PyMOL.

To gain a deeper understanding of the stability of protein-ligand docked complexes, MD simulation was carried out for the docked complexes of ligands targeting the substrate binding cleft and the S2-binding site, using GROMACS 2022.2 program with a CHARMM force field on Ubuntu-based LINUX workstation. For performing all MD simulations of protein-ligands, the topology files of selected ligands were generated using CHARMM36 General Force Field (CGenFF). The protein-ligand complexes were solvated in a cubic box and the steepest descent algorithm for 50,000 iteration steps was used to perform energy minimization. Two different system equilibration phases were carried out for 500,000 steps. The first phase of system equilibration was performed at a constant number of particles, volume, and temperature (NVT) at 300K with short range electrostatic cut-off of 1.2 nm. The second phase involved a constant number of particles, pressure, and temperature (NPT) ensemble at 300 K, for each step of 2 fs. Finally, the MD simulation was carried out for 100 ns with integration time frame steps of 2 fs. The RMSD values were calculated for PLpro protein and for all protein-ligand complexes.

### PLpro expression, purification, and inhibition assay

For expression in *E. coli*, the plasmid coding for PLpro was transformed into expression host *Rosetta* (DE3; Novagen, USA) and then cultured in 1 L LB broth medium at 37 °C. The expression of PLpro protein was induced using 0.5 mM isopropyl-β-d-1-thiogalactopyranoside (IPTG) along with the addition of 1mM ZnCl_2_ to log phase culture (OD_600_ = 0.6), and the cells were further grown overnight at 18 °C and 180 rpm in an incubator. Cells were harvested by centrifugation (6000 rpm for 10 min at 4 °C) and were resuspended in lysis buffer (50 mM Tris-HCl, pH 8.5, 500 mM NaCl, and 5mM DTT). Cells were homogenized using French press (Constant Systems Ltd, Daventry, England) and the cellular debris was removed by centrifugation at 10000 rpm for 1.5 h at 4 °C. The clarified supernatant was loaded onto pre-equilibrated nickel-nitrilotriacetic acid (Ni-NTA) (Qiagen, USA) affinity column and flow-through was collected after incubation for 30 min at 4 °C. After washing the resin with binding buffer (50 mM Tris-HCl, pH 8.5, 500 mM NaCl, and 5mM DTT), the recombinant histidine tagged PLpro protein was eluted with 100-500 mM imidazole in binding buffer. The eluted fractions were analyzed using 15% SDS-PAGE. The fractions containing pure PLpro protein were pooled together and dialyzed against buffer containing 20 mM Tris-HCl, pH 8.5, 200 mM NaCl, and 5mM DTT. The dialyzed protein was concentrated using Amicon centrifugal filters (Millipore, Burlington, MA, USA) with a molecular weight cutoff of 10 kDa. The concentration of purified PLpro was determined by using the Bradford assay. Expression and purification of C111S inactive PLpro mutant was performed using the same protocol as described for the wild type PLpro.

To characterize *in vitro* enzymatic activity of SARS-CoV-2 PLpro and to assess the inhibitory potential of identified compounds, a high-throughput FRET based enzymatic assay was developed as described previously (Choudhary et al., 2023; Shan et al., 2021). To carry this out, a commercially available fluorogenic substrate Z-Arg-Leu-Arg-Gly-Gly-7-amido-4-methylcoumarin (Z-RLRGG-AMC) (Biolinkk, India) representing the C-terminal residues of ubiquitin was used. The experiment was carried out in 96-well non-binding black plates (Corning) in reaction buffer containing 50 mM HEPES (pH 7.5). The protease diluted in reaction buffer was added to each well at a final concentration of 1 μM. Enzymatic reaction was initiated with the addition of fluorogenic substrate peptide at a concentration of 1.5 μM and the fluorescence signals were monitored continuously using a plate reader (Synergy HT, BioTek Instruments, Inc) with filters for excitation at 360/40 nm and emission at 460/40 nm for 30 min. PLpro C111S was used as a negative control to determine whether signal generation was dependent on proteolytic activity of enzyme. Data from duplicate set of readings was averaged together to plot a graph for relative fluorescence and time.

To assess the inhibitory potential of screened compounds, the assay was assembled as follows: 1 μM PLpro in 50 mM HEPES buffer was incubated with increased concentration of compounds (Cayman chemicals, USA) ranging from 0.1 μM to 500 μM and was dispensed into wells of 96-well plate. Reactions were initiated with 1.5 μM of fluorogenic substrate peptide and the fluorescence signals were monitored continuously for 10 min at excitation wavelength of 360/40 nm and emission wavelength of 460/40 nm. Data from duplicate set of readings were averaged together and a graph for percentage inhibition vs concentration was plotted for each selected compound after non-linear curve fitting using GraphPad Prism. Concentration of inhibitor that resulted in IC_50_ was calculated.

### Binding kinetic analysis using Surface Plasmon Resonance

The binding kinetics and affinity of identified compounds to PLpro of SARS-CoV-2 were analyzed by SPR using a Biacore T200 system (GE Healthcare) at 25 °C. In brief, histidine-tagged PLpro was diluted to 10 μΜ concentration at pH 7.3 and was then captured on NTA chip in 1X Phosphate buffer saline (1X PBS) running buffer (137 mM NaCl, 2.7 mM KCl, 10 mM Na_2_HPO_4_, and 1.8 mM KH_2_PO_4_) degassed and filtered through 0.22 μm membrane filter. The binding kinetic studies were performed by collecting the data at 25 °C by injecting the concentration series of analyte over the surface of chip having the immobilized PLpro with the following injection parameters: 10μL/min flow rate and 60 s association and dissociation phases. The surface was regenerated after each dissociation phase. Equilibrium dissociation constants (*K_D_*) of each pair of PLpro-ligand interaction were calculated using Biacore Evaluation Software (GE Healthcare) by fitting to a 1:1 Langmuir binding model.

### Viral growth and cytotoxicity assays in the presence of inhibitors

To determine the cytotoxic profiles of identified compounds on Vero cells, HEK293T-ACE2, and A549-ACE2, MTT assay was performed as described previously (Mudgal et al., 2022). Briefly, cells were seeded onto wells of 96-well cell-culture plate at a density of 1 × 10^4^ cells/well and the plates were incubated at 37 °C and 5% CO_2_ overnight for proper adherence of cells. The culture media was removed and the monolayer of cells was treated with increased dilutions of compounds (1-100 μM) prepared in DMEM media in triplicates. Post incubation period of 48 h, the media was removed, MTT [3-(4,5-dimethylthiazol-2-yl)-2,5-diphenyl tetrazolium bromide] solution (5 mg/mL) was added and incubated for 4 h at 37 °C in a humidified incubator with 5% CO_2_. Following this, the MTT solution was removed and the formazan crystals were dissolved in DMSO. Absorbance was measured at a wavelength of 550 nm using a multimode plate reader (Synergy HT, BioTek Instruments, Inc). Results were expressed as percentage viability of cells in comparison to untreated cell control. Percentage viability for each compound was calculated and a dose-response curve was plotted using GraphPad Prism software.

Viral load reduction assay was performed by quantitative reverse transcription-polymerase chain reaction (qRT-PCR). Briefly, Vero cells were seeded and cultured in 24-well plate in DMEM media supplemented with 10% FBS and 100 units of penicillin and 100 mg streptomycin/mL, at 37 °C and 5% CO_2_. 3 h before infection, the cells were washed with PBS and the media was replaced with DMEM (2% FBS) containing gradient of diluted experimental compounds. Post incubation, the cells were washed with PBS and were infected with SARS-CoV-2 at a Multiplicity of infection (MOI) of 0.01 in media without FBS and again incubated for 1.5 h. After incubation, the viral inoculum was removed and the cells were washed with PBS and then replenished with fresh medium containing dilutions of compounds. Plates were then incubated for 48 h at 37 °C and 5% CO_2_. At 48 hours post infection (hpi) the plates were freezed at −80°C. On the following day plate was thawed, and viral RNA was extracted from cell lysate of SARS-CoV-2 infected cells using HiPurA™ Viral RNA Purification Kit, according to the manufacturer’s instructions. One-step qRT-PCR was performed with primers targeting the SARS-CoV-2 envelope (E-gene) and RdRp region using the commercially available COVISure-COVID-19 Real-Time PCR kit (Genetix), as per the manufacturer’s instructions, to quantify the virus yield. The thermocycling conditions consist of 15 min at 50 °C for reverse transcription, 95 °C for 2 min, followed by 40 cycles of 30 s at 95 °C and 30 s at 60 °C. Azithromycin was used as positive control (data not shown). All experiments were performed in duplicates. Percentage inhibition was calculated based on ΔΔCt. The percentage inhibition versus concentration graph was plotted to calculate the EC_50_ values using the linear regression.

Infectious virus production by compound treated and infected cells as compared to only infected cells was also evaluated by the standard 50% tissue culture infectious dose (TCID_50_) and was expressed as TCID^50^/mL (Reed and Muench, 1938). For this purpose, 4 × 10^4^ Vero cells/well were seeded in a 96-well plate and the plate was incubated at 37 °C and 5 % CO_2_, till confluency. The freeze-thawed cell lysate of cells infected and treated with different concentrations of compounds collected at 48 hpi of the antiviral assay was subjected to 10-fold serial dilutions (10^1^ to 10^8^). Compound untreated and infected samples were used as positive control, and samples with only cell-culture media were used as a negative control. These dilutions were inoculated on Vero cells. Cells were incubated for 3 to 4 days at 37 °C and 5 % CO_2_. The plate was monitored daily for the appearance of a cytopathic effect (CPE). Post incubation, the viral inoculum was removed, and 10% formaldehyde (in PBS) was used to fix cells for 2 h at room temperature, followed by staining with 0.5% crystal violet solution for 1 h. Percent infected dilutions immediately above and immediately below 50% were determined. TCID_50_ was calculated on the basis of Reed and Muench method (Reed and Muench, 1938).

### Validation of antiviral activity in human cell lines

HEK293T-ACE2 and A549-ACE2 cells were seeded in 24 well plates and used for antiviral assay the next day. Single non-toxic concentrations of teneligliptin, linagliptin, lithocholic Acid or flupenthixol were prepared in infection medium (DMEM containing 2% FBS) and used to pre-treat cells for 3 h. For linagliptin and lithocholic acid, cells were treated with increasing doses of the drugs to estimate IC_50_. Cells were then washed with warm PBS and infected with 0.01 MOI SARS-CoV-2 in DMEM without FBS for 2 h. The inoculum was then removed, cells washed with PBS and incubated with infection media containing the drug concentrations. After 48 (HEK293T ACE2) or 72 h (A549 ACE2) total RNA was extracted using TRIzol (Thermo Fisher, 15596018) as per manufacturer’s instructions, and viral copy number was estimated by qRT PCR.

qRT-PCR: Equal amount of RNA was used to determine viral load using the AgPath-ID™ One-Step RT-PCR kit (Applied Biosystems, AM1005). The following primers and probes targeting the SARS-CoV-2 N-1 gene were used for amplification. Forward primer: 5’GACCCCAAAATCAGCGAAAT3’ and Reverse primer: 5’ TCTGGTTACTGCCAGTTGAATCTG3’, Probe: (6-FAM / BHQ-1) ACCCCGCATTACGTTTGGTGGACC. The Ct values were used to determine viral copy numbers by generating a standard curve using the SARS-CoV-2 genomic RNA standard. All statistical analyses were performed using GraphPad Prism 8.4.3 (GraphPad Software, USA). In all cases, a p-value < 0.05 is considered significant.

### Pro-inflammatory cytokine quantification and Interferon induction assay

HEK293T cells were seeded in the wells of 24-well plate 24 h prior to transfection. The media was removed and the cells were induced with 1 µg/mL polyinosinic:polycytidylic acid (poly I:C; Sigma) using Lipofectamine 2000 (Invitrogen). Post incubation period of 6 h, the media over the cells was removed, and fresh OptiMEM (Gibco) containing the compounds was added to cells (Devaraj et al., 2007; Thorne et al., 2022). 24 h post transfection, total RNA isolation was done using TRIzol (RNAiso; Takara) by following manufacturer’s instruction and was reverse-transcribed into first-strand cDNA using PrimeScript™ 1st strand cDNA Synthesis Kit (Takara Bio) in accordance with manufacturer’s protocol. For determination of effects of compounds on interferon, 500 ng of cloned pcDNA3 vector encoding PLpro protein of SARS-CoV-2 was transfected in HEK293T cells using Lipofectamine 2000 (Invitrogen) as per the manufacturer’s protocol. Cells were then stimulated with 1 µg/mL poly I:C (Sigma) using Lipofectamine 2000 (Invitrogen). After an incubation of 6 h, fresh OptiMEM media containing the compounds was added to cells. Mock treated cells or cells transfected with 500 ng of empty pcDNA3 vector or were included as controls (Devaraj et al., 2007; Shin et al., 2020; Thorne et al., 2022). RNA was isolated and was used for CDNA synthesis using PrimeScript 1st strand cDNA Synthesis Kit (Takara; India). The cDNA was used for Gene expression analyses for IL6, TNFα, and IFN-β cytokines, using KAPA SYBR fast universal qPCR kit (Sigma-Aldrich, USA) on QuantStudio real-time PCR system (Applied Biosystems, IN) with the expression of β-actin as the internal control. Based on the ΔΔCt method, The fold change in infected cells compared to corresponding controls was calculated after normalizing to the β-actin gene. All reactions were carried out in duplicates in a 96-well plate (Applied Biosystems, IN). Fold change was calculated by taking cells treated with only poly(I:C) as 1. The primer sequences used for qRT-PCR are listed in Table S3.

For interferon induction assay, HEK-293T cells were co-transfected with 50 ng IFNβ-luc firefly luciferase reporter plasmid, 20 ng RIGI 2CARD and 20 ng of pRL-TK Renilla luciferase reporter plasmid, along with/without 500 ng of plasmid expressing SARS-CoV-2 PLpro or empty vector. Transfection was done using Lipofectamine™ 2000 Transfection Reagent (11668019, Invitrogen) as per manufacturer’s instructions. 3 h post transfection, the media was replaced with complete medium containing 50 µM teneligliptin, 50 µM linagliptin, 25 µM lithocholic Acid or 10 µM flupenthixol. Cells were incubated for 24 h post transfection, then harvested in passive lysis buffer (E1941, Promega) and luciferase expression quantified using a dual luciferase assay (E1980, Promega). All statistical analyses were performed using GraphPad Prism 8.4.3 (GraphPad Software, USA). In all cases, a p-value < 0.05 is considered significant.

### Crystallization and structure determination

Purified SARS-CoV-2 PLpro inactive mutant form (C111S) was crystallized using sitting-drop vapor diffusion method. The purified protein was concentrated to ∼15 mg/mL and different protein-to-reservoir solution ratios (1:1, 2:1, and 1:2) were opted for crystallization with reservoir buffer containing crystallization condition [0.1 M Tris-HCl (pH 7.5), 1 M NaH2PO4, 1.2 M K2HPO4, and 10% Glycerol] in 96-well sitting-drop plate (Corning, USA). Diffraction quality crystals were produced after 1 week at 4 °C. The crystals were manually harvested from the drop and were cryo-protected by 10% glycerol and flash-frozen in liquid nitrogen for diffraction data collection. For co-crystallization of PLproC111S-lithocholic acid, a 5mM solution of lithocholic acid was added to purified PLproC111S (∼15 mg/mL) and the sample was incubated overnight at 4 °C. Further, the sample was used for setting up crystal trays using sitting drop vapor diffusion method at protein-to-reservoir ratio of 1:1 using crystallization condition similar to that for apo PLproC111S [0.1 M Tris-HCl (pH 7.5), 1 M NaH_2_PO_4_, 1.2 M K_2_HPO_4_ and 10% Glycerol]. For obtaining data of PLpro-linagliptin complex, co-crystallization was tried in the similar manner, but no crystals were produced. Therefore, soaking experiments were done to obtain PLpro-linagliptin complex data. The apo crystals for PLproC111S were transferred into crystallization solution supplemented with linagliptin (5 mM). The crystals were soaked for ∼6 h at 4 °C, and then these soaked crystals were cryoprotected using 10% glycerol.

X-ray diffraction data for PLproC111S apo and its complex with linagliptin and lithocholic acid were collected at the home source using Rigaku Micromax-007 HF high intensity microfocus rotating anode X-Ray generator equipped with Rigaku HyPix 6000C detector installed at Macromolecular Crystallography Unit, IIT Roorkee. Diffraction data were processed using the CrysAlis Pro software suite. Data reduction was performed using the Aimless program (Evans and Murshudov, 2013) from the CCP4i2 suite (Winn et al., 2011). The structures were solved by molecular replacement with MOLREP program (Vagin and Teplyakov, 2010) in CCP4i2 program suite using the structure of apo SARS-CoV-2 PLpro (PDB ID: 6WRH) as a reference model. The output model from molecular replacement was subsequently subjected to iterative cycles of model building using COOT (Emsley and Cowtan, 2004) and refinement using REFMAC (Murshudov et al., 2011). All structural analysis and figure preparation were carried out using PyMOL (Schrodinger LLC, 2015). The parameters for X-ray data collection and refinement statistics are presented in Table 2.

### HPLC

High-Performance Liquid Chromatography (HPLC) analysis for linagliptin was carried out on a Waters HPLC system equipped with Sunfire C18 reverse-phase column (250 × 4.6 mm, 5 µm particle size), 1525 binary pump and 2998 photodiode array detector (PDA). The degassed and vacuum filtered mobile phase consisted of water containing 0.1% Orthophosphoric acid pH 4.3 (A) and methanol (B) which were employed in the isocratic flow of 9A: 41B at a flow rate of 0.5 mL/min. Injection volume was set 20 µL and the column temperature was 28 °C. The ligand was monitored at 230 nm. Sample peaks were identified by matching retention time and UV spectrum with authentic standards.

### Animals and dosing

A total of 50 female mice, aged 6–8 weeks and weighing 18–20 g, were housed in a Biosafety Level-3 viral laboratory with unlimited access to feed and water. Linagliptin and molnupiravir were dissolved in 0.5% carboxymethyl cellulose (CMC) in PBS. Mice in the respective groups were either gavaged orally or injected intraperitoneally with 200 μL of the solution, starting 6 hours post-infection and continuing until sacrifice. The dosing regimen was 30 mg/kg or 60 mg/kg body weight (b.w.) for linagliptin, and 100 mg/kg or 200 mg/kg b.w. for molnupiravir. Control groups, both infected and uninfected, received 200 μL of 0.5% CMC alone (Figure 8). Animal studies and research protocols were approved by the Institute Animal Ethics Committee at Indian Institute of Science (IISc), Bangalore, India (IAEC approval no. CAF/Ethics/959/2020).

### Virus infection and animal sacrifice

The animals were anesthetized via intraperitoneal injection of ketamine (150 mg/kg, Bharat Parenterals Limited) and xylazine (10 mg/kg, Indian Immunologicals Ltd.), then intranasally challenged with 10⁵ plaque-forming units (PFU) of mouse-adapted SARS-CoV-2 (MA10) in 100 µL of PBS. Body weight was recorded daily, and the animals were sacrificed on day 4 post-infection using an overdose of xylazine. Lung samples were collected for virological analysis (right lobe) and histopathological examination (left lobe).

### Quantification of lung viral load by qRT PCR

Lung samples from each mouse were homogenized using a stainless-steel bead-based homogenizer, and total RNA was isolated following the manufacturer’s instructions for TRIzol (15596018, Thermo Fisher). Viral RNA was quantified using a 10µL reaction mixture containing 100ng of RNA per sample in a 384-well block, with the AgPath-ID™ One-Step RT-PCR kit (AM1005, Applied Biosystems). Primers and probes targeting the SARS-CoV-2 N-1 gene were used, with the following sequences: Forward primer: 5’ GACCCCAAAATCAGCGAAAT 3’, Reverse primer: 5’ TCTGGTTACTGCCAGTTGAATCTG 3’, and Probe: (6-FAM / BHQ-1) ACCCCGCATTACGTTTGGTGGACC. Ct values were used to determine viral copy numbers by generating a standard curve from a SARS-CoV-2 genomic RNA standard.

### Lung histology

Lung tissues were fixed in 4% paraformaldehyde in PBS and embedded in paraffin blocks. Tissue sections of 5 μm thickness were stained with Hematoxylin and Eosin (H&E). Slides were examined under light microscopy for three histological criteria in the lung (Alveolar infiltration and exudation, vasculature inflammation and peribronchiolar infiltration with epithelial desquamation) and were scored on a scale of 1 to 3 (mild=1, moderate=2, severe=3).

## Supporting information

supplementary file updated

## Acknowledgement

This research was funded by Science and Engineering Research Board, Department of Science & Technology, the Government of India (Proj. ref no. IPA/2020/000054). The authors acknowledge and thank Bioinformatics Center (BIC) supported by Department of Biotechnology, Govt of India (reference number BT/PR40141/BTIS/137/16/2021), Ashok Soota Molecular medicine facility, and the Macromolecular Crystallographic Unit (MCU) at Indian Institute of Technology, Roorkee. ST acknowledges funding support from BIRAC and Crypto Relief Fund. We thank the biosafety level-3 (BSL3) facility at Indian Institute of Technology, Roorkee (IIT Roorkee); Indian Institute of Science, Bengaluru, and Indian Veterinary Research Institute, Bareilly, India for support with the BSL3 facility and the procedures. S.C. thank the Council of Scientific & Industrial Research (CSIR), Government of India for financial support. S.N. acknowledges the Ministry of Human Resource Development, (MHRD). A.S thank Prime Minister’s Research Fellows (PMRF), Ministry of Education (MOE), Government of India for research fellowship.

## Data availability

All data are available in the main text or the supplementary materials. X-ray coordinates and structure factors are deposited at the RCSB Protein Data Bank under accession codes 8XTD and 8X1X.

