## supplementary file updated for "Activity profiling of deubiquitinating inhibitors-bound to SARS-CoV-2 papain like protease with antiviral efficacy in murine infection model"

***Table S1:*** *List of DUBs and cyanopyrrolidine ring inhibitors selected in in house drug library***.**

| **List of DUBs inhibitors screened in *in-house* library** | | | | | |
|---|---|---|---|---|---|
| • | Sitagliptin | • | E0AI3402 | • | WP1130 (Degrasny) |
| • | Alogliptin | • | HBX 19818 | • | GNE-6640 |
| • | Vildagliptin | • | HBX 91490 | • | GNE-7915 |
| • | Vildagliptin carboxylic acid | • | HBX 28364 | • | GNE0877 |
| • | Saxagliptin | • | HBX 41108 | • | AZ1 |
| • | Saxagliptin hydrate | • | HBX 28258 | • | AZ2 |
| • | 3-deoxy saxagliptin | • | Flupenthixol | • | AZ3 |
| • | Linaglitpin | • | PR619 | • | AZ4 |
| • | Anagliptin | • | P22077 | • | Cpd2 |
| • | Trelagliptin | • | P5091 | • | HY50736 |
| • | Gemigliptin | • | VLX1570 | • | HY5037A |
| • | Teneligliptin | • | b-AP15 | • | P22077 |
| • | Omarigliptin | • | Pimozide | • | P5091 |
| • | Evogliptin | • | GW7647 | • | MF-095 |
| • | Gasogliptin | • | RA-9 | • | Spautin-1 |
| • | Calpeptin | • | 6-Thioguanine | • | Mitoxantrone |
| • | Bisegliptin | • | Trifluoperazine | • | IU1-36 |
| • | Melogliptin | • | Rottlerin | • | IU1-47 |
| • | Denagliptin | • | SJB2-043 | • | IU1-248 |
| • | Retagliptin | • | SJB3-019A | • | auranofin |
| • | NVP DPP728 | • | ML323 | • | AZ1 |
| • | P32/98 | • | C527 | • | MF-094 |
| • | N-cyano pyrrolidine | • | 15-oxospiramilactone | • | Lithocholic Acid Hydroxyamide (LCAHA) |
| • | NSC632839 | • | ML364 | • | 2-hydroxy-piperidine |
| • | Curcusone D | • | XL188 | • | Lithocholic acid (LCAE) |
| • | NSC632839 | • | Compound 46 | • | FT671 |
| • | Compound 8QQ | • | AR946P0 | • | Vialidin A |

***Table S2:*** *List of top 20 hits identified after virtual screening of in house library along with binding energies.*

| Sr. No. | Name of compound | AutoDock Vina Binding energy (kcal/mol) |
|---|---|---|
| 1 | **Lithocholic acid hydroxyamide** | -8.3 |
| 2 | **HBX28364** | -7.9 |
| 3 | **Pimozide** | -7.8 |
| 4 | **Lithocholic acid (LCAE)** | -7.6 |
| 5 | **Gemigliptin** | -7.5 |
| 6 | **Linagliptin** | -7.5 |
| 7 | **MF094** | -7.5 |
| 8 | **FT671** | -7.4 |
| 9 | **ML323** | -7.4 |
| 10 | **XL188** | -7.3 |
| 11 | **Curcusone D** | -7.2 |
| 12 | **N-cyanopyrrolidine** | -7.2 |
| 13 | **Sitagliptin** | -7.2 |
| 14 | **HY50736** | -7 |
| 15 | **Denagliptin** | -6.9 |
| 16 | **Flupenthixol** | -6.9 |
| 17 | **FT827** | -6.9 |
| 18 | **GNE 6640** | -6.9 |
| 19 | **Teneligliptin** | -6.8 |
| 20 | **Vildagliptin** | -6.4 |

***Table S3:*** *Primer sequences used for qRT-PCR.*

| Gene | Forward primer | Reverse primer |
|---|---|---|
| IL6 | 5′-GCAGAAAAAGGCAAAGAATC-3′ | 5′-CTACATTTGCCGAAGAGC-3′ |
| IFN-β | 5′-TGGGAGGATTCTGCATTACC-3′ | 5′-AAGCAATTGTCCAGTCCCAG-3′ |
| CXCL10 | 5′-TGGCATTCAAGGAGTACCTC-3′ | 5′-TTGTAGCAATGATCTCAACACG-3′ |
| TNFα | 5′- AGCCTCTTCTCCTTCCTGATCGTG-3′ | 5′- GGCTGATTAGAGAGAGGTCCCTGG-3′ |
| β-actin | 5′-ATTGCCGACAGGATGCAGAA-3′ | 5′-GCTGATCCACATCTGCTGGAA-3' |
| PLpro_RT | 5′-TACCACCGATCCGAGCTTTCTG-3′ | 5′-CTGCCCATTTAATGCTGGTCAGAC-3' |

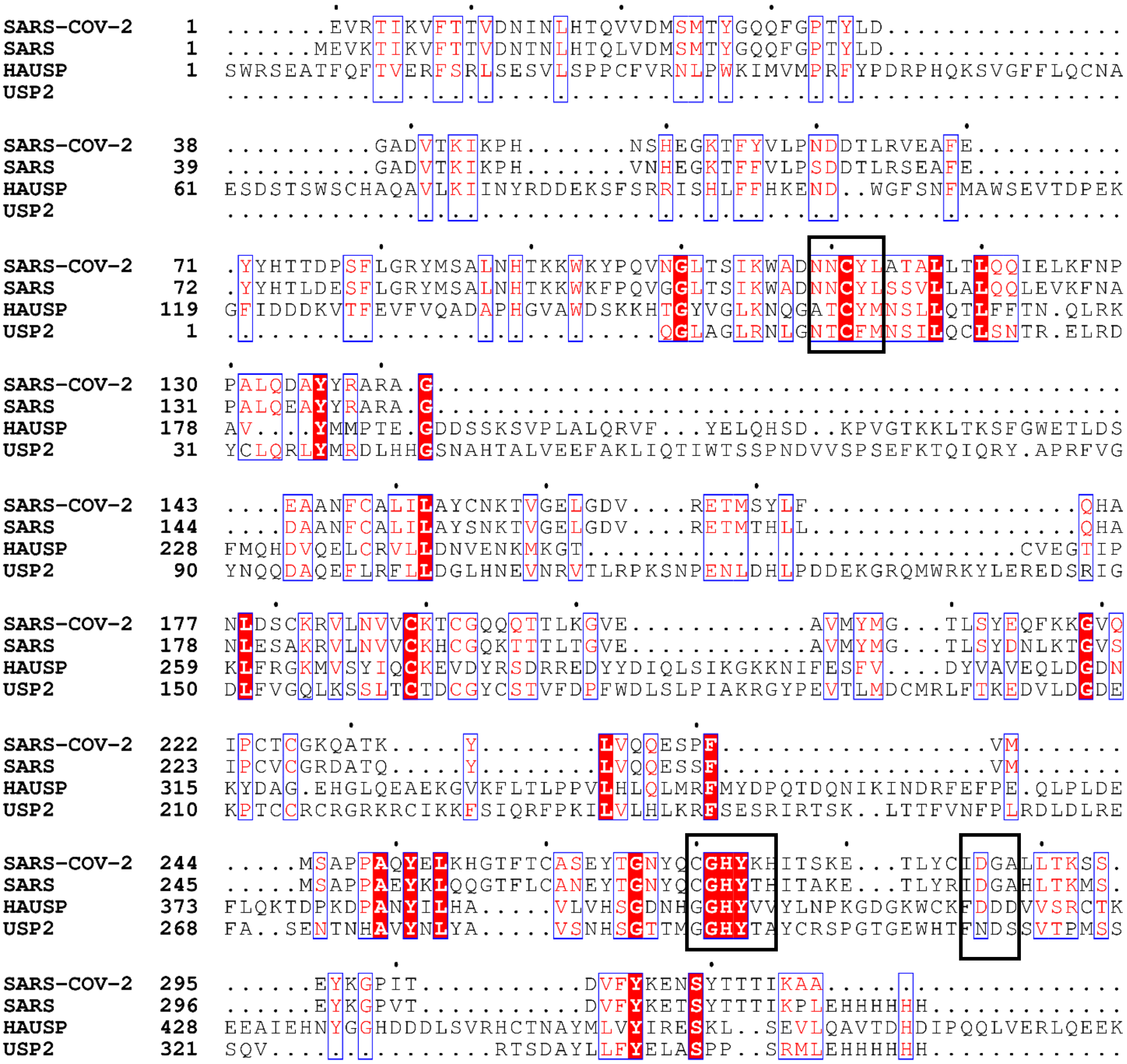


***Figure S1:*** *Sequence alignment for Deubiquitinating enzymes SARS-CoV-2 PLpro (PDB ID: 6W9C), SARS PLpro (PDB ID: 5E6J), USP2 (PDB ID: 5XU8), and USP14/HAUSP (PDB ID: 2F1Z). Multiple sequence alignment was generated using ESpript. Sequence numberings are represented according to PLpro of SARS-COV-2 (PDB ID: 6W9C). Conserved catalytic triad residues (Cys111, His 272, and Asp 286) are marked in boxes.*

**Substrate-binding cleft** **S2-Binding site**

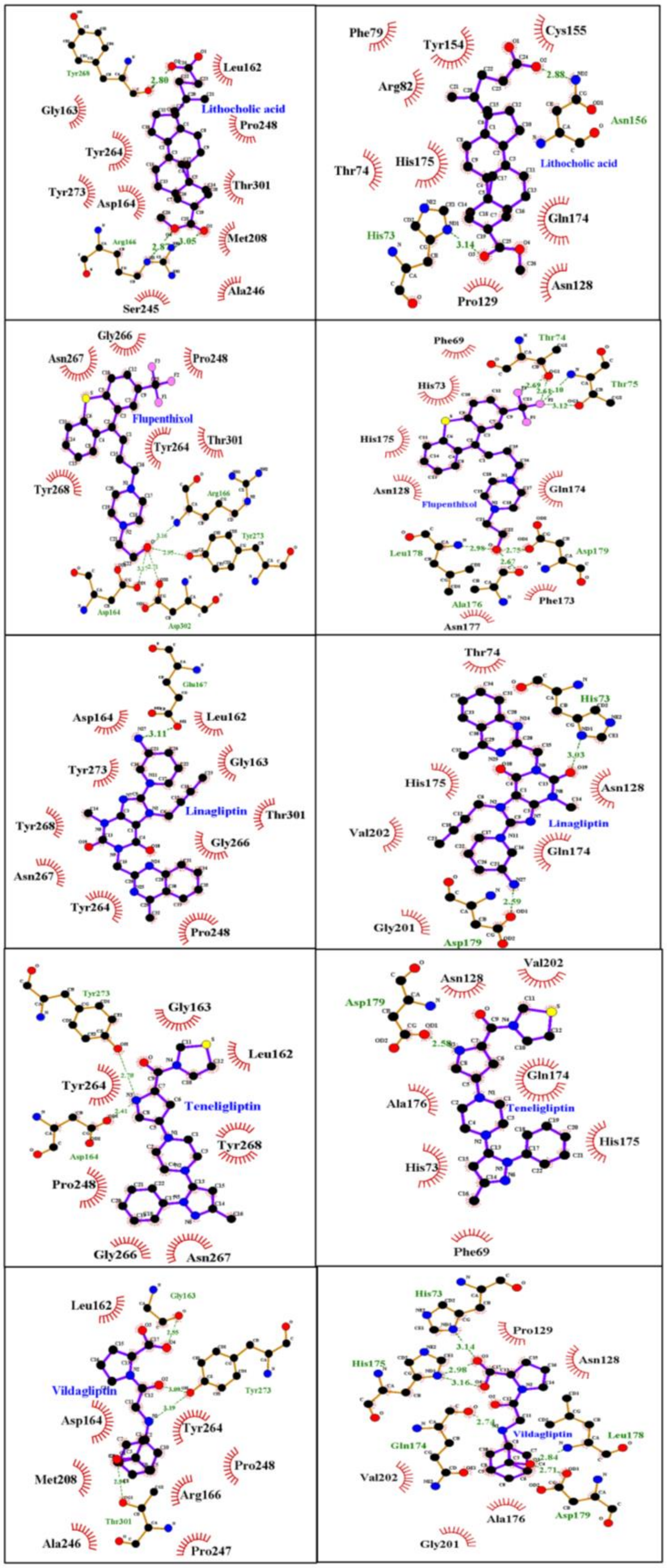


***Figure S2:*** *Two-dimensional interaction analysis of identified molecules targeting the substrate*

*binding cleft and S2-binding sites of PLpro*

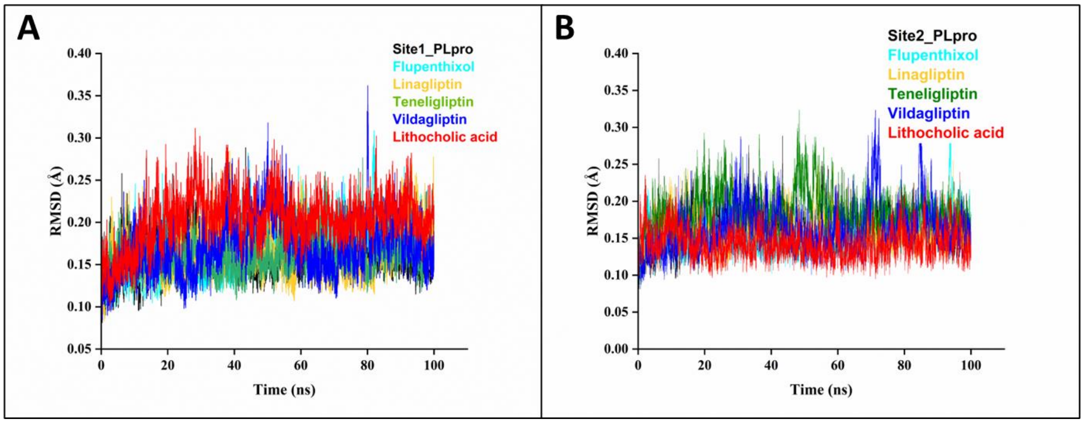


***Figure S3:*** *RMSD analysis of MD simulation trajectory. (a) RMSD plots obtained for PLpro in complex with Flupenthixol (cyan), Linagliptin (yellow), Teneligliptin (green), Lithocholic acid (red), and vildagliptin (blue) interacting at the substrate binding cleft (site1). (b) RMSD plots obtained for PLpro in complex with Flupenthixol (cyan), Linagliptin (yellow), Teneligliptin (green), Lithocholic acid (red), and vildagliptin (blue) interacting at the S2 site.*

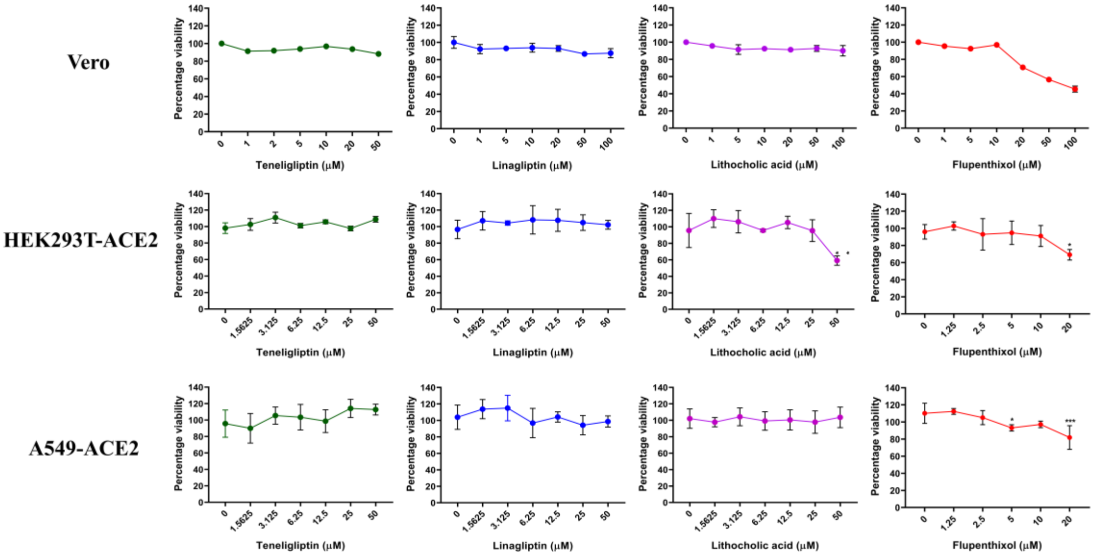


***Figure S4:*** *The cytotoxicity of Lithocholic acid, Linagliptin, Flupenthixol, and Teneligliptin in Vero, HEK293T-ACE2, and A549-ACE2 was determined by MTT assay. Cells were treated with increasing concentrations of Teneligliptin, Linagliptin, Lithocholic Acid or Flupenthixol as indicated, and after 48 h viability was estimated by MTT assay. Representative data are shown*

*from duplicate readings and the final graph was plotted for percentage viability of cells in the presence and absence of compounds (n = 2). *p < 0.05, **p < 0.01, ***p < 0.001; using One-way ANOVA with Dunnett's T3 multiple comparison tests. Error bars represent mean ± SD.*

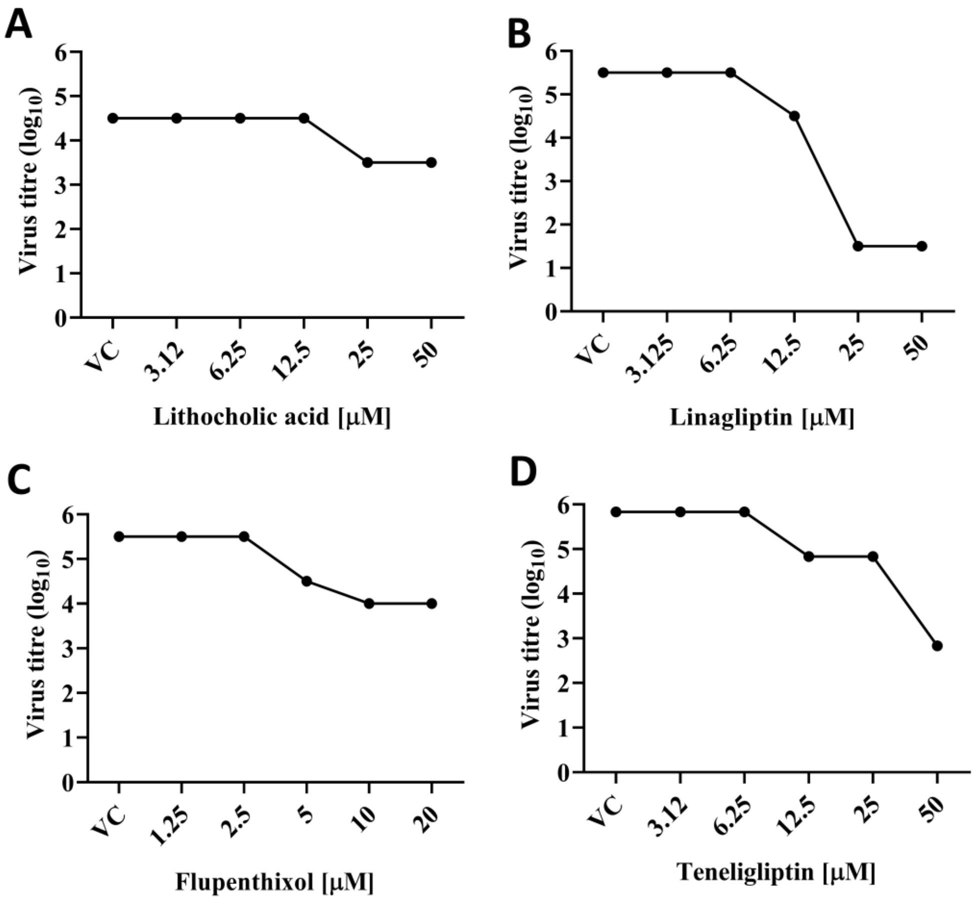


***Figure S5:*** *Graphs showing virus titer (TCID50/mL) of cell lysate of cells treated with increasing concentrations of compounds compared to virus control.*

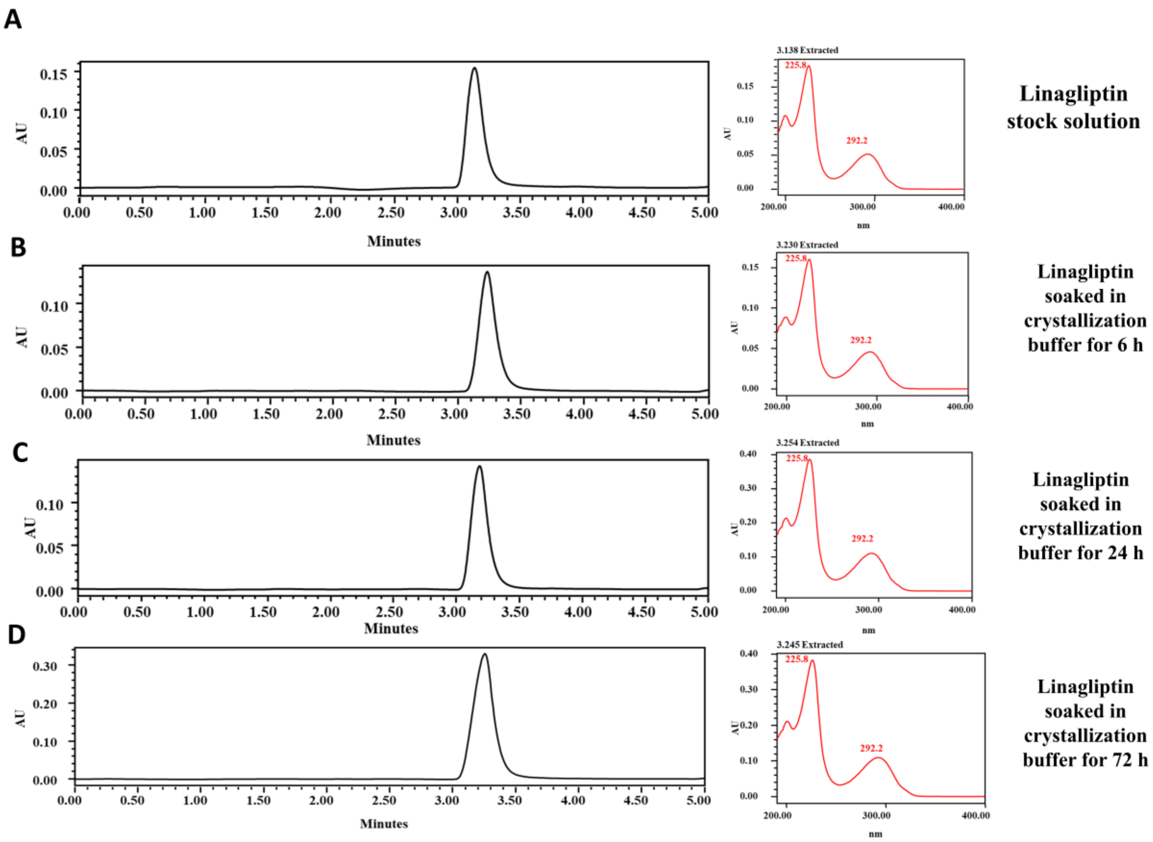


***Figure S6:*** *High Performance Liquid Chromatography (HPLC) chromatograms for (A) standard stock solution of Linagliptin, (B) Linagliptin soaked in crystallization buffer for 6 h, (C) Linagliptin soaked in crystallization buffer 24 h, and (D) Linagliptin soaked in crystallization buffer for 72 h.*
